# SCAD Delivery Platform: A Novel Approach for Efficient CNS and Extrahepatic Oligonucleotide Therapeutics

**DOI:** 10.1101/2024.10.29.620792

**Authors:** Moorim Kang, Wei-Hsiang Lin, Feifei Xu, Xiaojie Zhou, Chunyan Si, Kunshan Liu, Jianxiu Dai, Jichao He, Ian Schacht, Yichen Li, Zubao Gan, Long-Cheng Li

## Abstract

Oligonucleotide therapeutics, including antisense oligonucleotides (ASOs) and duplex RNAs like siRNA, saRNA, and miRNA, hold immense potential for treating genetic and acquired diseases by modulating gene expression in a target-specific manner. However, effective delivery to extrahepatic tissues, particularly the central nervous system (CNS), remains a significant challenge. While GalNAc conjugation has enabled liver-specific delivery, leading to several approved siRNA drugs for hepatic targets, CNS delivery lags. ASOs, on the other hand, can self-deliver to the CNS when administered locally, as seen with nusinersen and tofersen. To address this disparity, we’ve developed the Smart Chemistry Aided Delivery (SCAD) platform which enables duplex RNA delivery by conjugating it to an accessory oligonucleotide (ACO), which acts as an aptamer to mediate protein binding and facilitate cellular uptake. Through extensive screening, we identified an optimal SCAD architecture that demonstrates enhanced cell-free protein binding and *in vitro* activity. In rodent models, local administration of SCAD-siRNA conjugates resulted in broad biodistribution throughout the CNS and sustained mRNA knockdown for over five months, with a favorable safety profile. The SCAD platform also exhibited efficient delivery to other tissues, including the eye, the lung and the joint. These features support its potential for broader clinical applications, as evidenced by an ongoing trial targeting amyotrophic lateral sclerosis (ALS) associated with mutations in the *SOD1* gene. The modular design of SCAD allows it to easily adapt to any duplex RNA, making it a powerful tool for advancing oligonucleotide therapeutics.

## Introduction

Oligonucleotide-based therapies have emerged as powerful tools for targeting both genetic and acquired diseases at the molecular level. Antisense oligonucleotides (ASOs) and duplex RNAs— including siRNA, saRNA, and miRNA—modulate gene expression at the transcriptional and post-transcriptional levels in a target specific manner, providing avenues to treat conditions that are challenging to address with traditional small molecules or biologics (Ranasinghe et al., 2023). However, delivering these molecules to target tissues, particularly beyond the liver, presents significant challenges (Gupta et al., 2021, Tang and Khvorova, 2024). While the liver can be effectively targeted using technologies such as GalNAc conjugation, achieving similar delivery efficiency to other tissues, especially the central nervous system (CNS), has remained elusive (D’Souza et al., 2023). Various strategies have been explored to overcome these barriers, including lipid nanoparticles, chemical conjugations (such as sugar, lipophilic, peptide and protein moieties), ligand-targeted nanoparticles, all aimed at targeting specific tissues like the eye, lung, and CNS (Roberts et al., 2020, Brown et al., 2022, Wickline et al., 2023).

Antisense oligonucleotides (ASOs) have demonstrated effectiveness for neurological disorders, exemplified by the FDA approval of nusinersen for spinal muscular atrophy (SMA) and tofersen for amyotrophic lateral sclerosis (ALS) associated with mutations in the *SOD1* gene (Crooke et al., 2021, Blair, 2023). These successes demonstrate the potential for localized CNS delivery (Hill and Meisler, 2021, Amanat et al., 2022). Although ASOs and siRNAs both regulate gene expression, they significantly differ in their delivery mechanisms and molecular structures. Both are inherently negatively charged, which typically impedes cellular uptake. However, phosphorothioate (PS) modifications substantially enhance the distribution and cellular uptake of ASOs and improve their overall biological activity by increasing nuclease resistance, stability, and binding affinity to plasma proteins like albumin and lipoproteins, which facilitates their cellular entry (Eckstein, 2014, Crooke et al., 2017, Crooke et al., 2021, McCampbell et al., 2018, Crooke et al., 2020). Conversely, in siRNAs, extensive use of PS modifications can reduce their therapeutic activity by interfering with RNA-induced silencing complex (RISC) loading and increase toxicity, thus limiting their applicability in similar contexts (Alterman et al., 2019).

To overcome the challenges associated with delivering duplex RNA into CNS, we developed the Smart Chemistry Aided Delivery (SCAD) platform. This innovative system utilizes the self-delivery properties of ASOs by conjugating a non-targeting single-stranded oligonucleotide, known as an accessory oligonucleotide (ACO), to the duplex RNA. Our prior published studies have demonstrated the effectiveness of SCAD in delivering a siRNA targeting *SOD1* (si*SOD1*) to the CNS. Local delivery of si*SOD1*-ACO conjugate via intracerebroventricular (ICV) or intrathecal (IT) injection into *SOD1^G93A^*mice slowed disease progression, extended animal survival, and mitigated disease-related motor deficits (Duan et al., 2024). In non-human primates, intrathecal administration of si*SOD1*-ACO conjugate led to a robust and durable dose-dependent knockdown of *SOD1* mRNA in the CNS and a decrease in SOD1 protein in the cerebrospinal fluid (CSF) (Duan et al., 2023). These encouraging results have led to the initiation of an investigator-initiated trial (IIT, NCT05903690) and a phase I study (NCT06556394) to evaluate the safety and tolerability of RAG-17, a SCAD-siRNA conjugate, in ALS patients with *SOD1* mutations.

The present work outlines the development of SCAD technology and presents detailed pharmacodynamic and pharmacokinetic analyses of SCAD after local administration in the CNS. Additionally, we have investigated SCAD’s capacity to target the lung, the eye and the joint, highlighting its potential for broader therapeutic applications. Collectively, our findings underscore SCAD’s potential as a transformative delivery platform for oligonucleotide therapeutics, offering delivery solutions to target gene modulation across a broad spectrum of diseases.

## Results

### Structure and synthesis of the SCAD architecture

The SCAD delivery system is structured with a single-stranded oligonucleotide, termed an accessory oligonucleotide (ACO), which is linked via a linker to one of the strands (the passenger strand) in a double-stranded RNA (dsRNA). The passenger strand, the linker and the ACO were synthesized on solid support as a single strand and annealed with the other strand of the dsRNA to form the final SCAD architecture (Fig. 1A). The initial ACO was designed to have no significant homology with any human coding sequences and consisted of 18 nucleotides (nt) featuring a universal phosphorothioate (PS) backbone and 2’-O-methoxyethyl (2’-MOE) modifications.

**Figure 1.**
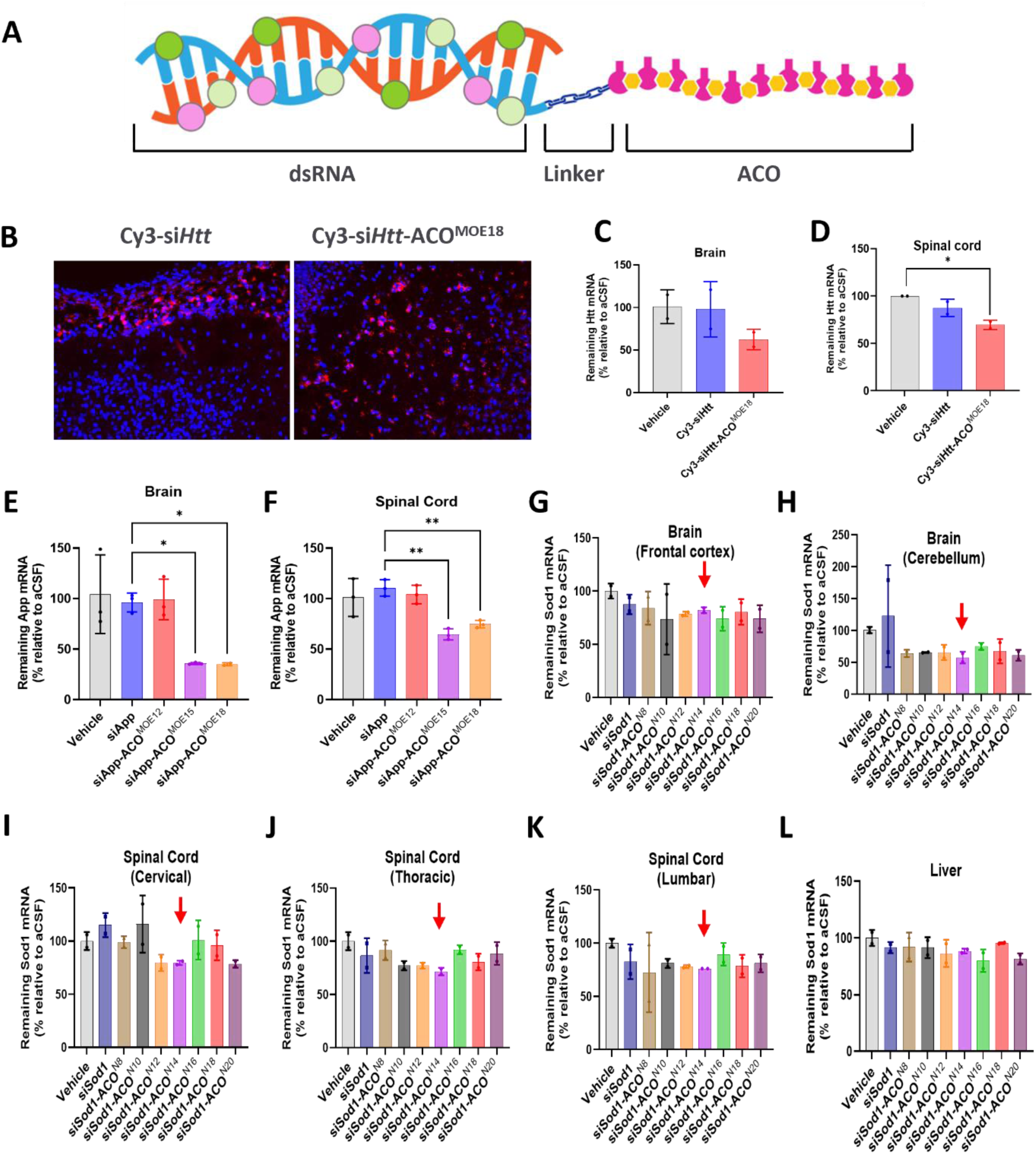
Schematic of the SCAD architecture and *in vivo* proof-of-concept testing. **A**. The SCAD architecture consisting of a dsRNA conjugated to an accessory oligo (ACO) via a linker. **B-D**. SCAD enhanced siRNA distribution and activity in mouse CNS. C57BL/6J pup mice (PND 3-5, 2.5 g) were administrated a single dose of 0.1 mg of either Cy3-siHtt (final concentration 7.6 nmol) or Cy3-si*Htt*-ACO2^MOE18^ (final concentration 4.8 nmol) via ICV injection. After 3 days the brain was harvested and sectioned. The sections were counterstained with DAPI before fluorescent microscopy (B). Mouse *Htt* expression in brain (C) and spinal cord (D) was quantified by RT-qPCR. **E** and **F**. The effect of ACO size on siRNA activity *in vivo*. C57BL/6J pup mice (PND 3-5, 2.5 g) were given a 0.1 mg ICV injection of si*App* or SCAD variants (si*App*-ACO^MOE12^, si*App*-ACO^MOE15^, and si*App*-ACO^MOE18^) with final concentrations of 7.6, 5.5, 5.1, and 4.8 nmol respectively. After 3 days, brain and spinal cord *App* expression was measured by RT-qPCR. **G-K**. C57BL/6J mice (4-6 weeks) were given a 0.1 mg ICV injection of si*Sod1* or SCAD variants (si*Sod1*-ACO^N8^, si*Sod1*-ACO^N10^, si*Sod1*-ACO^N12^, si*Sod1*-ACO^N14^, si*Sod1*-ACO^N16^, si*Sod1*-ACO^N18^, and si*Sod1*-ACO^N20^) with final concentrations of 7.2, 6.2, 5.9, 5.7, 5.5, 5.3, 5.1, and 4.9 nmol respectively. After 5 days, *Sod1* expression was measured by RT-qPCR in the brain and spinal cord. Statistical significance was determined by *one-way* ANOVA and *Dunnett’s* multiple comparisons test. *\*, p < 0.05; **, p < 0.01*.

The concept was initially applied to a Cy3 labeled duplex siRNA targeting mouse *Htt* gene to create si*Htt*-ACO^MOE18^ which was administered via ICV to neonatal mice. For comparison, the same Cy3-labeled siRNA without the ACO was also administered as a control. Three days post-administration, si*Htt*-ACO^MOE18^ showed widespread distribution across various brain regions, whereas the control siRNA’s fluorescence was confined to the vicinity of the injection site in the lateral ventricle (Fig. 1B). Consistent with these observations, si*Htt*-ACO^MOE18^ achieved significant reduction in *Htt* mRNA levels in both the brain and spinal cord, unlike the control siRNA which had no significant effect on mRNA levels (Fig. 1C and 1D).

To further validate the SCAD concept, the approach was applied to another siRNA for mouse *App* gene (siApp). ACOs of varying lengths (12, 15, and 18 nt) were conjugated to si*App*. The 15 and 18 nt ACO variants (si*App*-ACO^MOE15^, si*App*-ACO^MOE18^) effectively knocked down *App* expression in the brain and spinal cord, whereas the 12 nt ACO (si*App*-ACO^MOE12^) did not (Fig. 1E and 1F). These results indicate that SCAD is broadly applicable to siRNAs and that ACO size matters for delivery efficiency.

In the si*Htt*-ACO^MOE18^ study intended for a longer observational duration, all mice succumbed by day 6 post-dosing, indicating potential toxicity associated with the ACO’s chemical makeup. In response, we developed a range of ACOs, varying from 20-nt to 8-nt, based on the original sequence but omitting the 2’-MOE modifications across all nucleotides. These variants were conjugated to a siRNA targeting the mouse *Sod1* gene and administered via ICV to older mice aged 4-6 weeks. None of the mice displayed mortality before the planned sacrifice on day 5. Subsequent tissue analysis showed that these 2’-MOE-lacking variants exhibited diminished knockdown efficiency in the CNS (Fig. 1G-1K) when compared to their 2’-MOE counterparts (Fig. 1E and 1F). However, among all the variants, the 14-nt ACO variant stood out as the most effective variant across most of the tissues (Fig. 1G-1K). This finding emphasizes the impact of ACO size and chemical properties on the biodistribution and effectiveness of SCAD, underscoring the need for further optimization of the SCAD architecture.

### Optimization of the SCAD architecture and *in vitro* evaluation

Having established the benefits of ACO conjugation in delivering duplex RNA *in vivo,* we developed an *in vitro* screening method aimed at rapidly evaluating a large library of SCAD designs. Drawing from the established benefits of chemical modifications in ASOs, such as the phosphorothioate (PS) backbone and 2’-O modifications (2’-O-methyl or 2’-O-methoxyethyl), which are known to enhance plasma protein binding and facilitate cellular uptake(Crooke et al., 2020), we initiated our study by establishing an *in vitro* protein binding assay.

In this assay, the duplex RNA si*Sod1* conjugated to a 14-nt ACO with uniform 2’MOE modification (si*Sod1*-ACO^MOE14^) or without the ACO was incubated with human plasma for 1 hour at 37°C, followed by electrophoretic mobility shift assays (EMSA). The results, as depicted in Figure 2A, showed that si*Sod1*-ACO^MOE14^ had a significantly lower unbound fraction compared to its unconjugated counterpart, indicating enhanced plasma protein binding attributable to the ACO. This finding supports the notion that ACO conjugation enhances plasma protein interaction.

**Figure 2.**
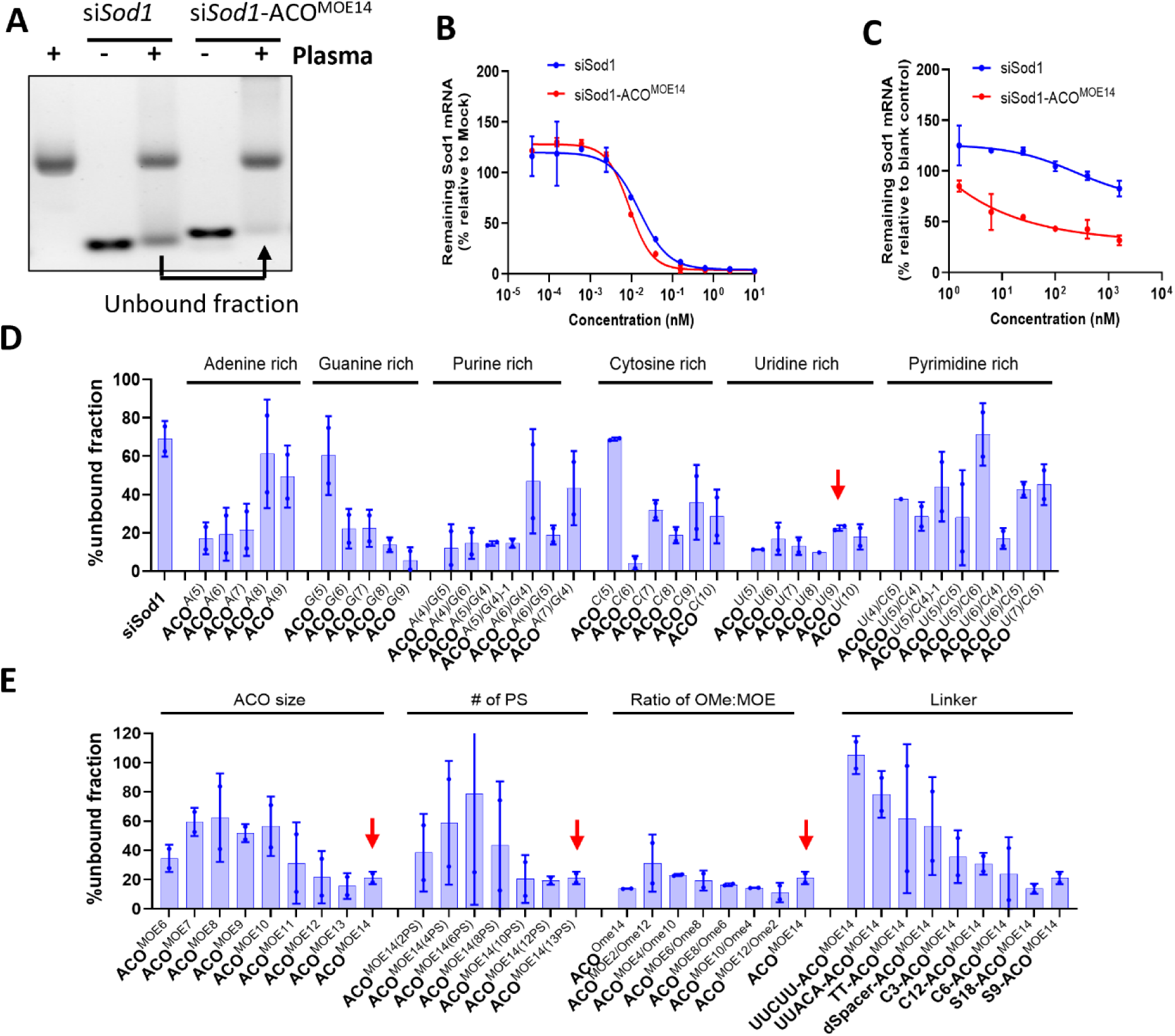
SCAD optimization with protein binding assay. **A.** Protein binding of siSod1 and siSod1-ACO^MOE14^ was assessed by EMSA after incubation with mouse plasma for 1 hour at 37°C. **B** and **C**. Knockdown activity of siSod1 and siSod1-ACO^MOE14^ (or ACO^MOE14^) in primary mouse hepatocytes (PMH) transfected by RNAiMax for 24 h **(B)** or free uptake for 3 days **(C)**. Sod1 expression was analyzed by RT-qPCR, normalized to a mock or blank control (PMH without oligonucleotides). **D** and **E**. Protein binding of different SCAD designs was assessed by EMSA as in (A) and quantified by gel dosimetry. Values on y axis are unbound fraction (means ± SEM of two replicates samples). The red downward arrow indicates the best ACO sequence identified by the screen.

We further developed an *in vitro* assay to assess SCAD’s delivery efficacy using unassisted delivery mechanisms in freshly isolated primary mouse hepatocytes (PMHs). Initially, we compared the knockdown efficacy of siRNA with or without the SCAD architecture, both delivered using the RNAiMAX transfection reagent. As shown in Figure 2B, there was no significant difference in the knockdown of Sod1 expression between si*Sod1* and si*Sod1*-ACO^MOE14^, suggesting that ACO conjugation does not compromise the *in vitro* potency of the siRNA.

Subsequently, we directly administered both si*Sod1* and si*Sod1*-ACO^MOE14^ to the PMHs without any transfection aid. Twenty-four hours later, Sod1 mRNA expression was measured using RT-qPCR. The results revealed that si*Sod1*-ACO^MOE14^ exhibited significantly stronger knockdown of Sod1 expression compared to the non-conjugated siRNA (Fig. 2C). This outcome validates the effectiveness of the *in vitro* free uptake assay for evaluating the delivery performance of SCAD-conjugated siRNAs.

### SCAD library development and functional analysis

To optimize the SCAD system further, we developed an extensive library of ACO sequences varying in sequence context, nucleotide composition, size, and 2’ sugar modifications. This included creating ACOs enriched with different nucleotide bases—adenine, guanine, purine, cytosine, uridine, and pyrimidine—maintained at a constant length of 14 nucleotides with full 2’-MOE and thirteen PS backbone modifications. We also varied the lengths of these ACOs from 6 to 14 nucleotides, maintaining full 2’-MOE and PS backbone modifications. We then adjusted the PS content in these 14-nt ACOs, ranging from 2 to 13 PS modifications. Additionally, we experimented with different ratios of 2’-MOE to 2’-OMe in ACOs of a fixed 14-nt length with full PS backbones. Finally, we explored the use of two types of linkers: flat linear spacers (C3, C12, C6, S9, S18) and more rigid, ring-shaped cyclic spacers (TT dinucleotide, dspacer, UUACA, UUCUU), to connect the 14-nt ACO with universal 2’-MOE and PS backbone modifications to the AS strand of si*Sod1*.

We first assessed the protein binding capabilities of these SCAD variants. As shown in Figure 2D and 2E, discernible trends emerged in the impact of various modifications on protein binding. For instance, protein binding decreased with increased adenine and purine content and increased with higher guanine content, larger ACO sizes, or greater numbers of PS modifications. However, changes in the content of purine, pyrimidine, and the ratio of 2’-MOE to 2’-OMe showed negligible effects on protein binding. Notably, linear spacers facilitated stronger protein binding compared to cyclic spacers.

Next, we evaluated the *in vitro* knockdown activity of these SCAD variants in PMHs using the free uptake approach. Results, as shown in Figure 3A and 3B, indicated that the nucleotide composition significantly affected activity; for example, increasing adenine or adenine and guanine content enhanced activity, while an increase in cytosine content reduced it. Among different linkers, TT and S9 provided the siRNA with the highest knockdown activity (Fig. 3B).

**Figure 3.**
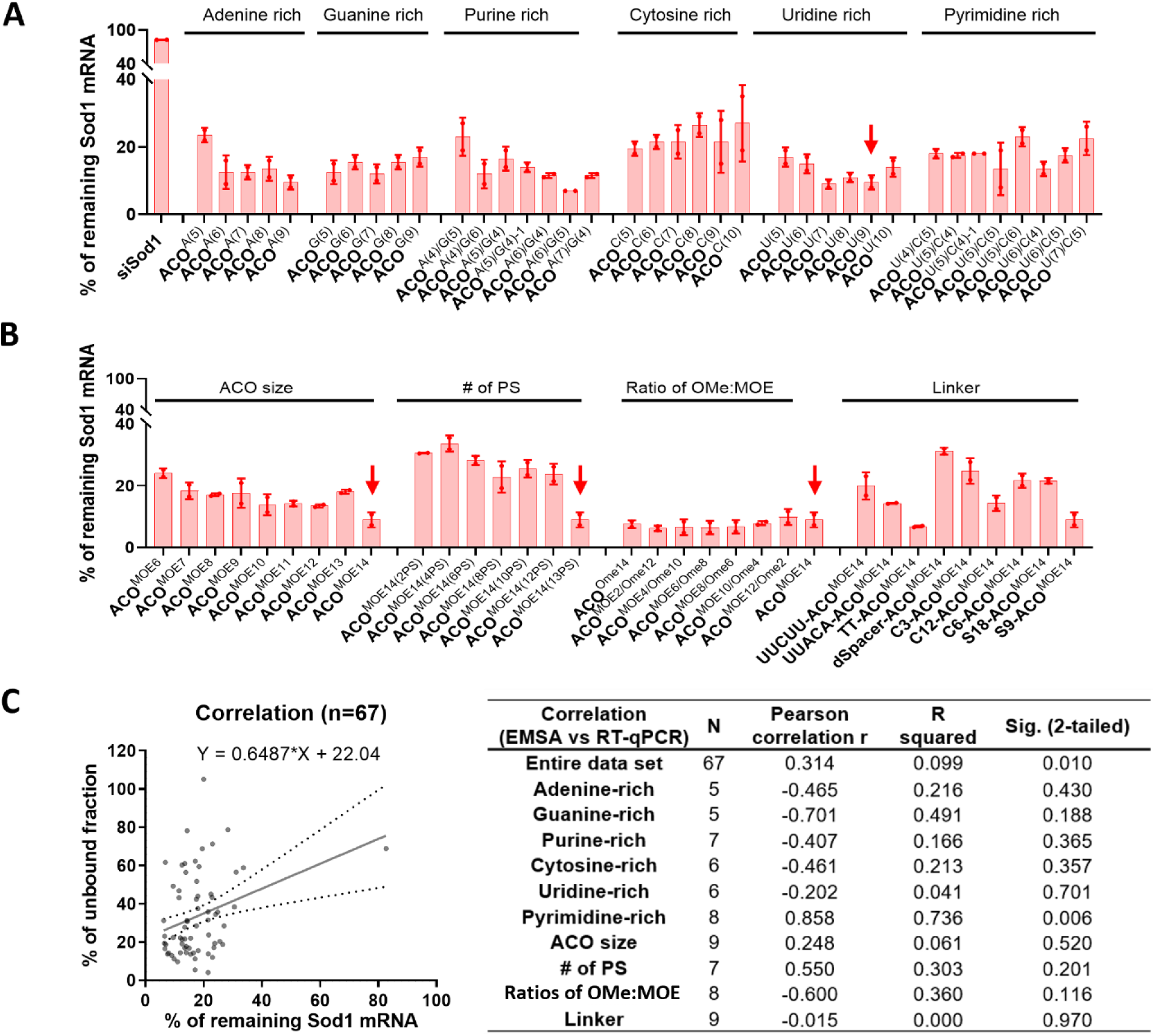
SCAD optimization with *in vitro* knockdown activity. **A** and **B**. The indicated SCAD variants were directly added to PMH at 1 mM and 24 h later the treated cells were harvested for mRNA expression assessment by RT-qPCR. Data represents means ± SD of two replicate treatments and is plotted as % of remaining Sod1 mRNA relative to untreated cells after normalizing to Tbp and Hprt1 gene. The red downward arrow indicates the best ACO sequence identified by the screen. **C**. Correlation of protein binding ability (EMSA data, % of unbound fraction) with knockdown activity (RT-qPCR data). Sig, statistical significance.

We then performed correlation analysis between the protein binding ability and knockdown activity for each variant. As shown in Figure 3C, there was almost no correlation (R squared = 0.099) between protein binding and knockdown activity when all variants were combined. When the correlation analysis was separately done for each group of variants, we could see that pyrimidine-rich ACOs gave the strongest correlation (R squared = 0.736) between protein binding and activity, followed by guanine-rich ACOs (R squared = 0.491). PS content and ratio of 2’-Ome:2’-MOE in the ACOs showed weak correlation between protein binding and activity (R squared = 0.303 and 0.36 respectively).

Together, these findings indicate that the careful selection and optimization of ACO characteristics—particularly concerning nucleotide compositions and linker types—are crucial for enhancing the SCAD system’s efficacy.

### *in vivo* validation of knockdown activity in the CNS

To determine if the *in vitro* findings could be replicated *in vivo*, we administered selected SCAD candidates to mice via ICV injection and assessed *Sod1* mRNA expression in CNS tissues 7 days post-injection. We included optimal candidates, as well as suboptimal candidates as controls. The optimal candidates all have strong protein binding, which include the following designs: purine-rich (ACO^A(6)/G(4)^), fully 2’-Ome modified (ACO^Ome14^), uridine-rich (ACO^U(7)^ and ACO^U(9)^), ACO^MOE14^ with spacer 9 (S9-ACO^MOE14^), with 6-carbon spacer (C6-ACO^MOE14^), or with spacer 18 (S18-ACO^MOE14^). The suboptimal candidates have designs that include spacer 9 without ACO (S9), cytosine-rich (ACO^C(10)^), 14-nt ACO with two PS backbones (ACO^MOE14(2PS)^), and ACO with a TT linker (TT-ACO^MOE14^) which has poor protein binding.

As depicted in Figure 4, optimal candidates (ACO^A(6)/G(4)^, ACO^Ome14^, ACO^U(7)^, ACO^MOE14^, S9-ACO^MOE14^, C6-ACO^MOE14^, S18-ACO^MOE14^) in general displayed stronger knockdown activities especially in the spinal cord and the liver comparing to suboptimal candidates (S9, ACO^C(10)^, ACO^MOE14(2PS)^, TT-ACO^MOE14^). Notably, reducing the number of PS in the ACO (ACO^MOE14(2PS)^) almost completely abolished knockdown activity. Among the optimal candidates, uridine-rich ones (ACO^U(7)^ and ACO^MOE14^) demonstrated substantial knockdown activities in the brain and the spinal cord. Furthermore, ACO^MOE14^ with different linkers exhibited robust knockdown activities (38.9%, 72.9%, 77.5%, and 76.8%) in the cerebrum (Fig. 4A).

**Figure 4.**
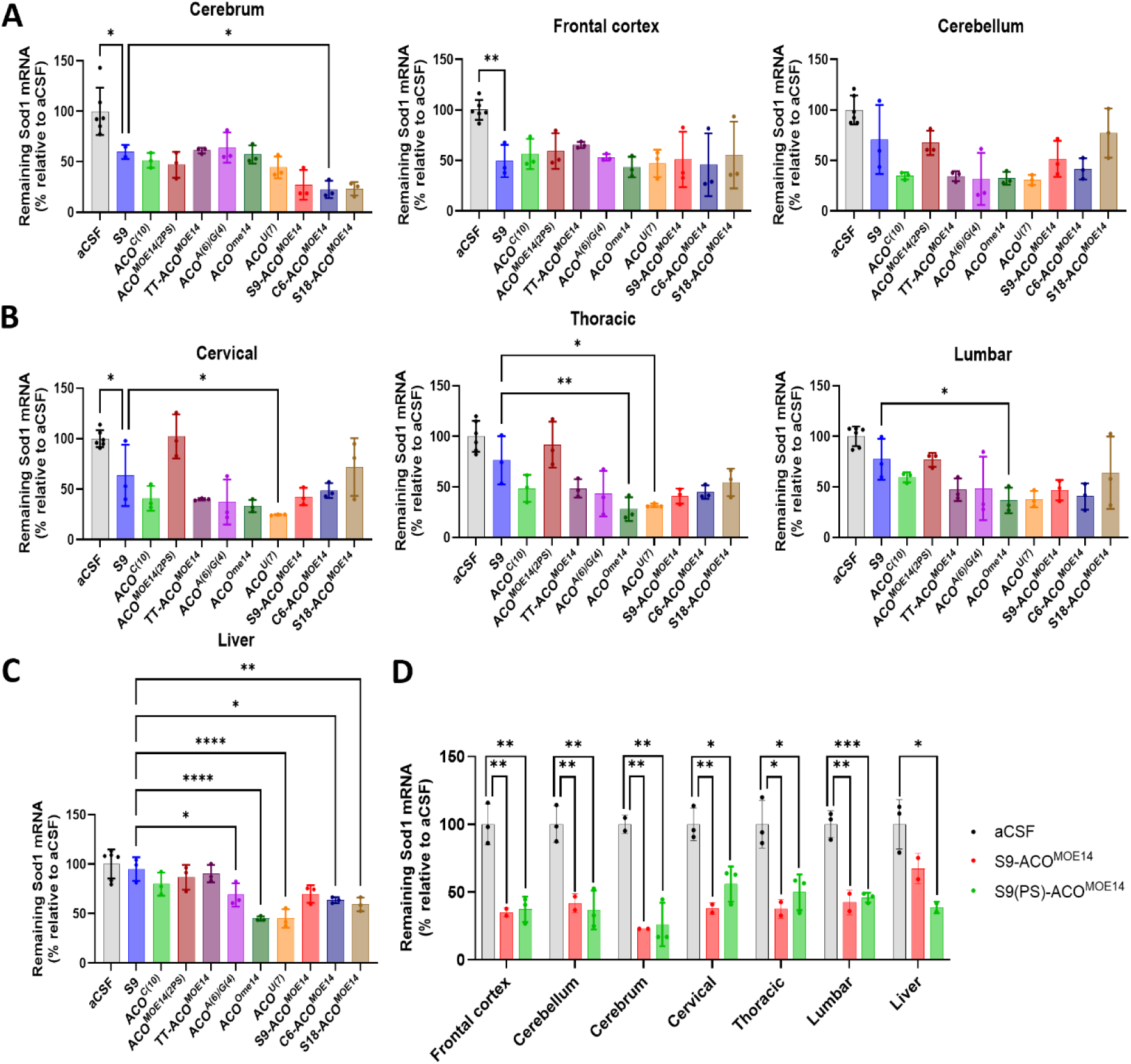
Validating effective SCAD designs in mice. C57BL/6J mice were administrated a single dose of 0.2 mg ICV injection of si*Sod1*, with either spacer 9 without ACO (S9) or the indicated SCAD variants (ACO^C(10)^, ACO^MOE14(2PS)^, TT-ACO^MOE14^, ACO^A(6)G(4)^, ACO^Ome14^, ACO^U(7)^, S9-ACO^MOE14^, C6-ACO^MOE14^, and S18-ACO^MOE14^) at a final concentrations of at 14.3, 10.2, 10.3, 10.0, 10.2, 10.6, 10.2, 10.2, 10.2, and 10.1 nmol respectively. The vehicle control group received aCSF. Mouse Sod1 expression was quantified by RT-qPCR using gene-specific primers in tissues from different regions: **(A)** Brain (*i.e.*, cerebrum, frontal cortex, cerebellum), **(B)** Spinal cord (*i.e.*, cervical, thoracic, lumbar), or **(C)** Liver on day 7 after treatment. (**D**) C57BL/6J mice were administrated a single ICV dose of 0.2 mg siSod1, with either cleavable linker (S9-ACO^MOE14^) or uncleavable linker (S9(PS)-ACO^MOE14^). Sod1 mRNA expression was quantified as in A-C 7 days postdosing. Statistical significance was determined by *one-way* ANOVA and *Dunnett’s* multiple comparisons test. *\*, p < 0.05; **, p < 0.01; ****, p < 0.0001*.

These results validate that the combination of cell-free protein binding assays and *in vitro* free uptake activity serves as an effective preliminary screening method for identifying efficient SCAD architectures for *in vivo* delivery. The findings from this *in vivo* study established the optimized ACO design, particularly ACO^MOE14^ with the S9 linker (S9-ACO^MOE14^), as a standard for effective siRNA delivery to the CNS with limited systemic exposure, setting the stage for further development of the SCAD system.

We reasoned that the stability of the S9-ACO^MOE14^ design could be compromised *in vivo* due to the phosphodiester (PO) linkages at both sides of the S9 linker. To address this potential issue, we replaced the PO linkages with phosphorothioate (PS) bonds, resulting in a new variant, S9(PS)-ACO^MOE14^. Unfortunately, this alteration led to reduced efficacy in the spinal cord and unintended increased activity in the liver on day 7 post-ICV administration (Fig. 4D). This outcome suggests that the additional PS modifications may exacerbate CNS drainage, reducing activity in the spinal cord while increasing the unwanted liver activity.

### SCAD enabled broad CNS tissue distribution, and profound and durable siRNA activity in the CNS

To evaluate long-term durability and PD/PK relationship of CNS delivered SCAD-siRNA, a single dose of SCAD-s*iSod1* or duplex si*Sod1* was administered to mice via ICV injection and tissues were harvested at different time points (1 and 8 hours and 1, 3, 14, 30, 61, 92 and 154 days post-injection) to assess siRNA tissue concentration by stem-loop PCR (all time points) and *Sod1* mRNA expression by RT-qPCR (1, 3, 14, 30, 61, 92 and 154 days).

The results, detailed in Figure 5, Figure S1, and Table S1, reveal that SCAD-si*Sod1* exhibited a broader and more sustained distribution in CNS tissues compared to duplex si*Sod1*. Notably, SCAD-si*Sod1* concentrations remained higher than those of si*Sod1* over extended period of time in the frontal cortex, motor cortex, and hippocampus, indicating improved retention. The ratio of SCAD-si*Sod1*’s concentration to si*Sod1*’s was particularly high in the hypothalamus, brain stem, and cerebellum, with peak ratios observed at later time points (Table S1), suggesting sustained retention facilitated by SCAD. The spinal cord showed modest exposure levels of SCAD-si*Sod1*, which could be attributed to its anatomical distance from the ICV injection site.

**Figure 5.**
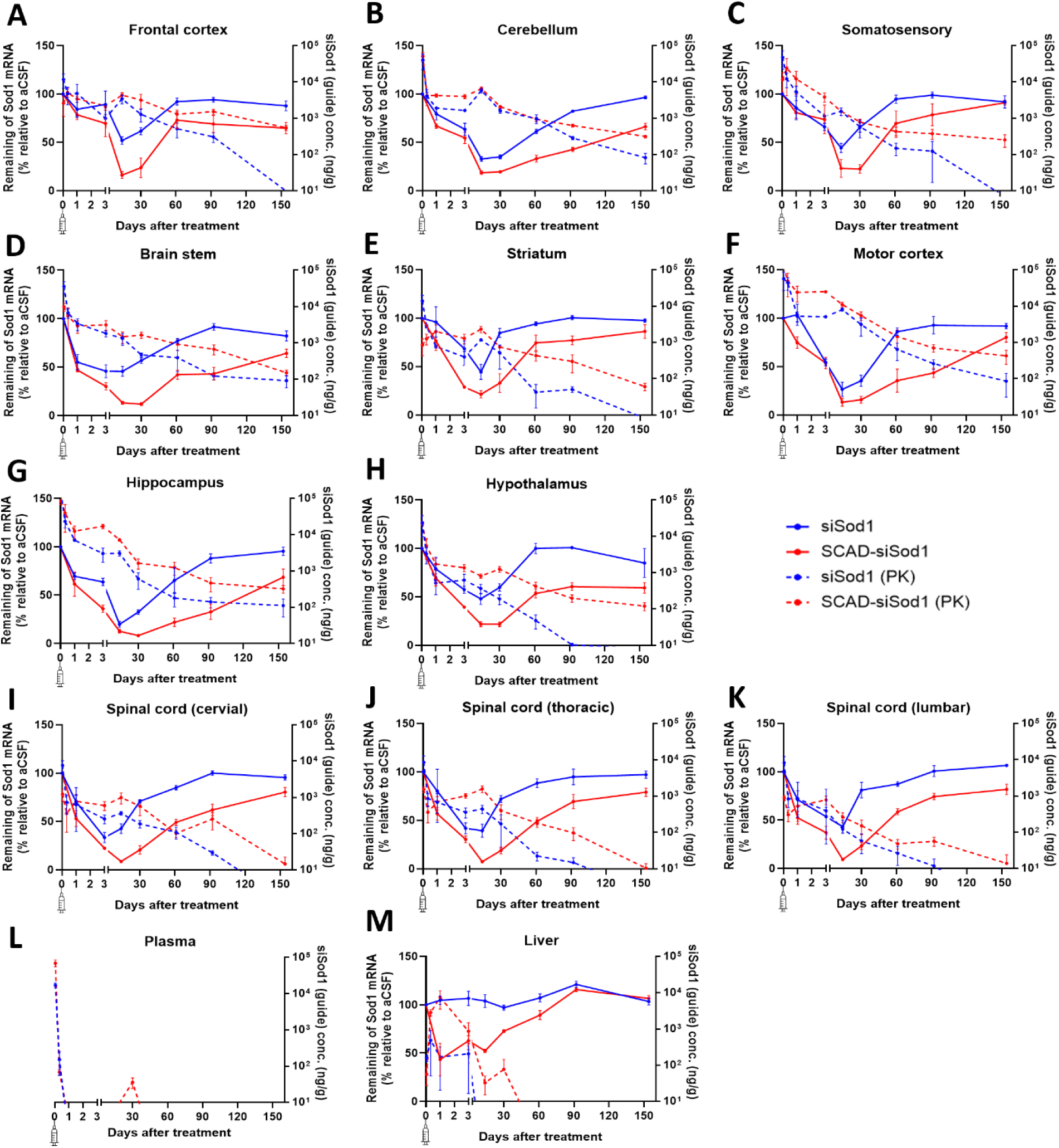
SCAD-siRNA achieved broad tissue exposure and demonstrated durable knockdown activity in mouse CNS. C57BL/6J mice were administrated with either siSod1 or SCAD-si*Sod1* at a fixed molecular dose of 21 nmol via ICV injection or received aCSF as a vehicle control. The concentration of siRNA (specifically, the guide strand) and the expression of *Sod1* were analyzed in various tissues at different time points over a 154-day period at 1h, 8h, as well as on day 1, 3, 14, 30, 61, 92 and 154 after dosing using stem-loop RT-qPCR and RT-qPCR respectively. The expression of mouse Sod1 and the concentration of siRNA in the brain (*i.e.*, frontal cortex (**A**), cerebellum (**B**), somatosensory (**C**), brain stem (**D**), striatum (**E**), motor cortex (**F**), hippocampus (**G**), and hypothalamus (**H**)), as well as the spinal cord (*i.e.*, cervical (**I**), thoracic (**J**), and lumbar (**K**)), were analyzed. Peripheral tissues (*i.e.*, plasma (**L**), and liver (**M)** were also analyzed. However, Sod1 expression was not analyzed at 1h and 8h after dosing, as well as in the plasma samples. The data represent the mean tissue concentration and expression values ±SD, with n=3-4 mice per group. The dotted lines in the graph indicate 50% or 80% knockdown.

In the plasma, both si*Sod1* and SCAD-si*Sod1* showed initial peak concentration at 1-hour post-dosing, and then rapidly declined to undetectable levels within one day, indicating quick systemic clearance (Fig. 5L, Table S1). In the kidney and liver, si*Sod1* and SCAD-si*Sod1* reached peak concentration at 8 hours and 1 day respectively, suggesting quicker clearance of si*Sod1* than SCAD-si*Sod1* (Fig. 5M, Table S1).

The PD assessments showed that, in contrast to the duplex siRNA, SCAD-si*Sod1* induced profound knockdown effects across all CNS regions, including deep brain structures such as the brain stem, striatum, hippocampus, and hypothalamus. Peak knockdown activities approached or exceeded 80% between days 14 and 30 (Fig. 5, Fig. S1). In comparison, si*Sod1* alone achieved notable knockdown only in regions close to the ICV injection site—the hippocampus and the motor cortex—with effectiveness peaking at 40.4% and 64.7%, respectively.

Remarkably, while the knockdown effects of the duplex siRNA diminished rapidly, SCAD-si*Sod1* maintained substantial activity, with over 30% knockdown persisting in the frontal cortex, cerebellum, brainstem, hippocampus, and hypothalamus as late as day 154 (Fig. 5A-K, Fig. S1). This sustained activity illustrates the robust PD/PK relationship across most CNS tissues, with IC50 values ranging from 0.47 nmol in the lumbar spinal cord to 24 nmol in the somatosensory cortex (Fig. S2), confirming that tissue siRNA concentration reliably predicts *in vivo* activity.

Overall, these findings demonstrate that SCAD delivery via ICV injection achieves efficient and sustained siRNA delivery to the CNS, resulting in prolonged tissue exposure and persistent pharmacodynamic activity for over five months.

### Cell-type specificity of SCAD-siRNA in the CNS

To assess the cell-type-specific activity of SCAD in the CNS, we synthesized siRNAs targeting genes specifically expressed in mouse astrocytes, microglia, and neurons—namely *Gfap*, *Iba1*, and *Tubb3*. Additionally, we targeted the rat neuron-specific gene, *Map2*, as previously documented (Brown et al., 2022). These siRNAs, featuring uniform chemical modifications and the SCAD architecture, were administered to mice via a single ICV injection and to rats via a single intrathecal (IT) injection and the expression of these target genes was analyzed 14 days after administration.

In mouse CNS, all SCAD-siRNAs effectively reduced the mRNA levels of their respective targets, with the most significant reduction observed in astrocytes, followed by neurons and microglia (Fig. 6A-C). For rats, the neuron-specific siRNA targeting *Map2* displayed the most profound activity in the spinal cord, with significant but reduced activity throughout most brain regions (Fig. 6D). This gradient of activity, from the highest in the spinal cord to lower levels in the brain could be explained by the fact that intrathecally injected SCAD-siRNA moved counter to the direction of CSF flow, unlike the ICV-injected SCAD-siRNA in mice which moved with the CSF flow from the brain to the spinal cord.

**Figure 6.**
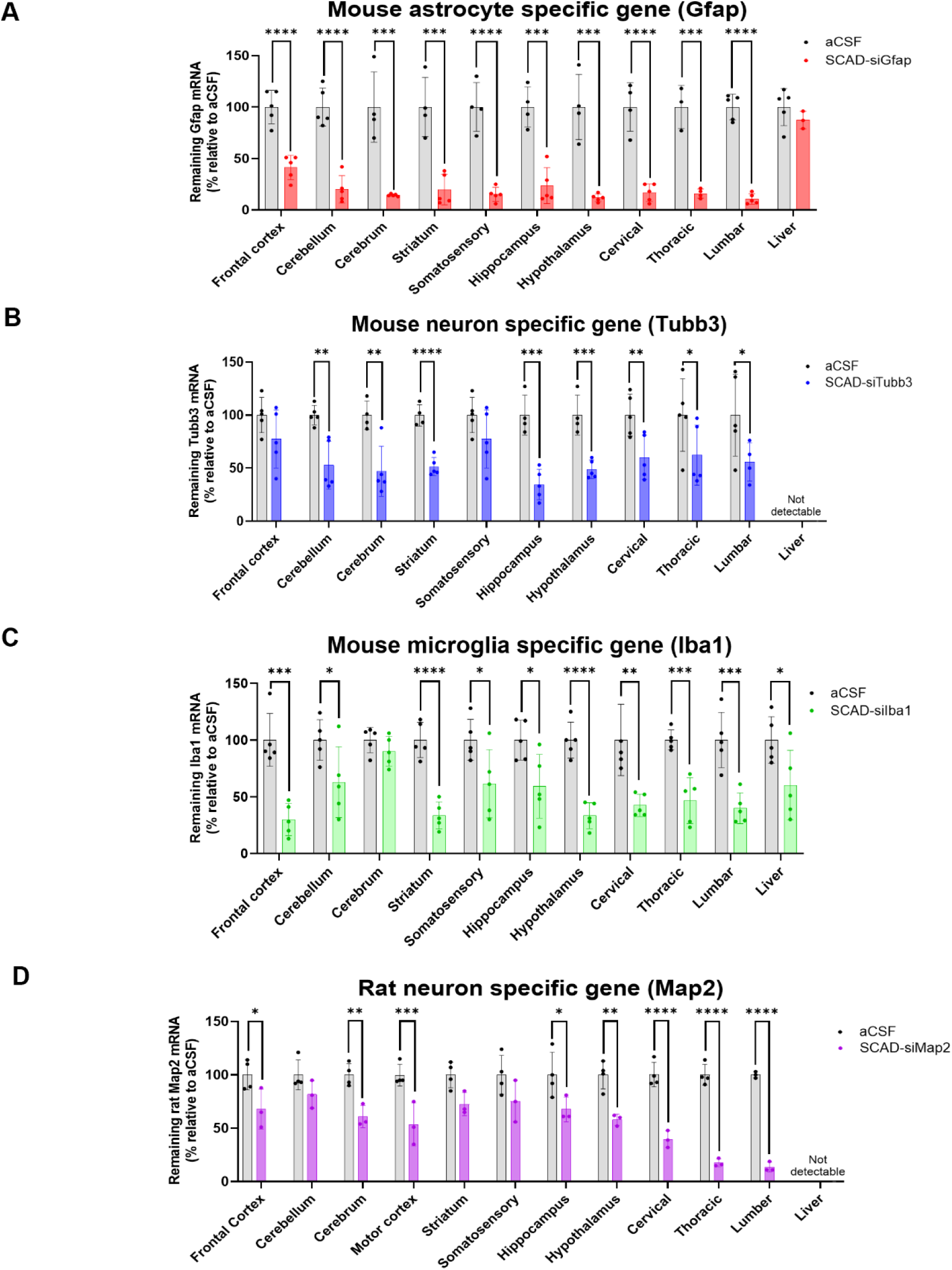
Cell type specificity of SCAD-siRNA in mouse CNS. C57BL/6J received a single dose (0.2 mg) of SCAD-siRNAs that targeted **(A)** the astrocyte-specific gene *Gfap*, **(B)** the neuron-specific gene *Tubb3*, and **(C)** the microglia-specific gene *Iba1* via ICV injection. (**D**) SD rats received a single dose (0.9 mg) of SCAD-si*Map2* targeting neuron specific gene *Map2* via IT injection. After 14 days, the brain, spinal cord, and liver were collected to assess siRNA knockdown by RT-qPCR. It was found that *Tubb3* genes were not detectable in liver tissue. The data presented are the mean expression values ±SD (n = 4-5 animal per group) of each target gene relative to aCSF vehicle treatment. Statistical significance was determined by an unpaired *t-* test (one-tailed). *\*, P < 0.05; **, P < 0.01; ***, P < 0.001; ****, P < 0.0001*.

Comparing knockdown efficacy across these cell types requires consideration of the intrinsic potency of each siRNA and the baseline expression levels of the target genes. Despite these variables, the results clearly demonstrate that SCAD can effectively deliver siRNA across a broad range of cell types in the CNS, confirming its significant efficacy.

### SCAD delivered duplex RNA to additional extrahepatic tissues (lung, eye, and joint) via local administration

#### Lung delivery by intratracheal instillation

To assess the efficacy of SCAD for lung delivery, si*Sod1* and SCAD-si*Sod1* were administered to mice via intratracheal instillation. *Sod1* expression was evaluated 7 days post-administration. While siSod1 alone led to a slight decrease in *Sod1* mRNA expression, SCAD-si*Sod1* achieved statistically significant, dose-dependent reductions in *Sod1* levels (p < 0.001 or p < 0.0001), with even the lowest dose of 1 nmol resulting in a 43.6% reduction (Fig. 7A).

**Figure 7.**
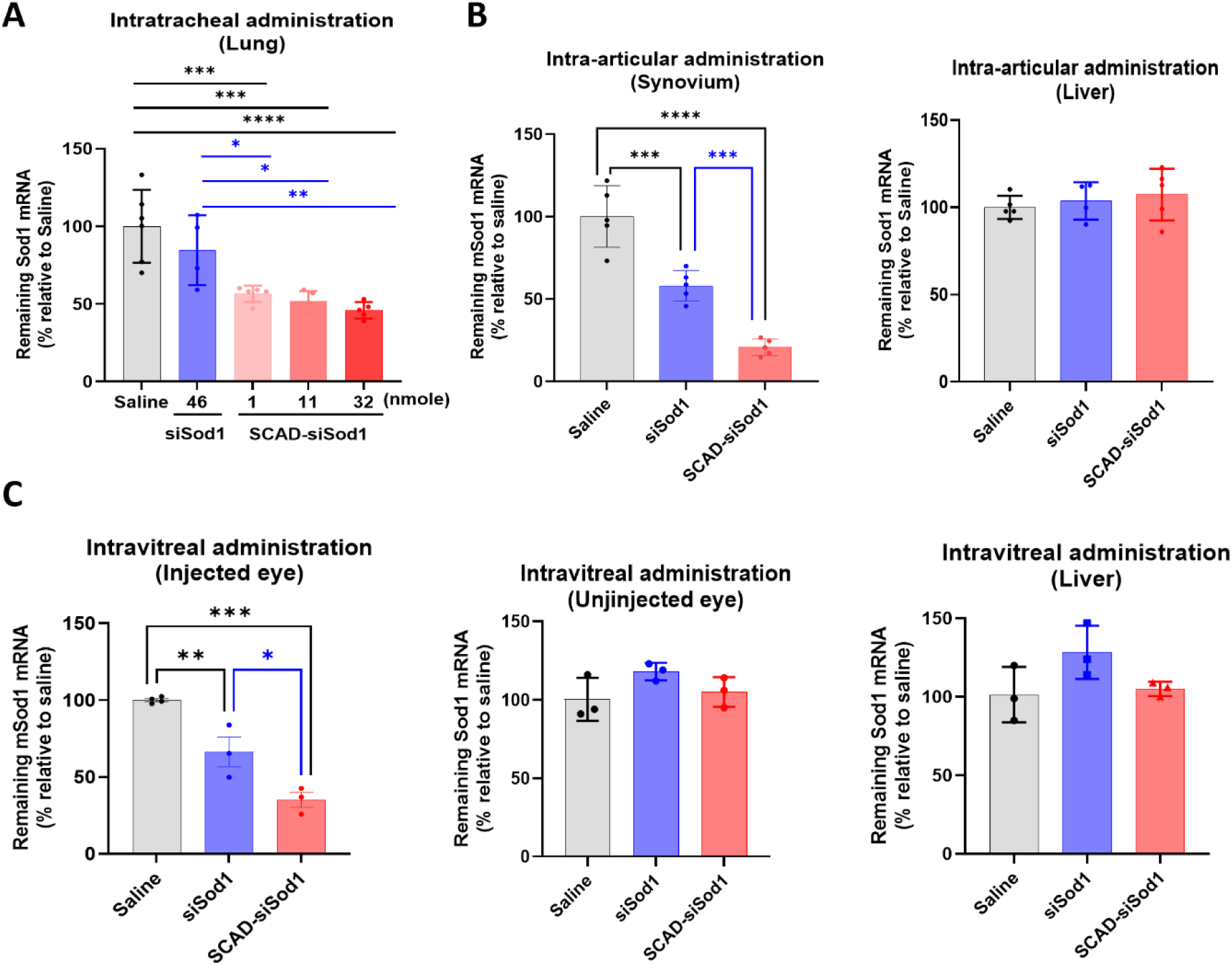
SCAD delivered duplex RNA into extrahepatic tissues, including the lung, eye and joint. **A.** C57BL/6J mice received either si*Sod1* or SCAD-si*Sod1* at the indicated doses via intratracheal injection (n = 4-6 mice per group). The mice were sacrificed 7 days after dosing. **B.** SD rats received either si*Sod1* or SCAD-si*Sod1* via intra-articular injection at a fixed molecular dose of 7 nmol (n = 3-4 rat eyes per group). The rats were sacrificed 14 days after dosing. **C.** SD rats received a single dose of 30 mg per eye of either si*Sod1* (final concentration 2.1 nmol) or SCAD-si*Sod1* (final concentration 1.6 nmol) via intravitreal injection. The rats were sacrificed 14 days after dosing. The expression of *Sod1* in mouse or rat tissues from the lung, retina, synovium or liver was quantified via RT-qPCR. Saline alone was used as the vehicle control to establish baseline expression. The data presented are the mean expression values ±SD of the target gene relative to saline treatment. Statistical significance was determined by *one-way* ANOVA and *Dunnett’s* multiple comparisons test. *\*, P < 0.05; **, P < 0.01; ***, P < 0.001; ****, P < 0.0001*.

#### Joint delivery by intraarticular injection

The ability of SCAD to target joint tissues was examined in rats that received 7 nmol of si*Sod1* or SCAD-si*Sod1* via intraarticular injection. Fourteen days later, analysis of the synovial membrane revealed that SCAD-siSod1 led to a 79.3% reduction in *Sod1* expression, significantly outperforming the 41.9% reduction observed with si*Sod1* alone (Fig. 7B). Notably, neither form of the siRNA showed detectable activity in the liver, confirming localized action within the joint.

#### Eye delivery by intravitreal injection

To evaluate the potential of SCAD for eye delivery, si*Sod1* and SCAD-si*Sod1* were administered intravitreally at doses of 2.1 and 1.6 nmol, respectively, to rats. Fourteen days later, *Sod1* mRNA levels in retina tissues were assessed. SCAD-si*Sod1* achieved a 64.9% reduction in *Sod1* expression, markedly surpassing the 33.7% reduction seen with si*Sod1* (Fig. 7C). Crucially, there were no changes in *Sod1* mRNA levels in the un-injected eyes or liver, indicating minimal systemic exposure.

These results collectively confirm SCAD’s capability to efficiently deliver RNA therapeutics to a variety of extrahepatic tissues, achieving significant and localized gene silencing with minimal systemic involvement.

### Safety and toxicological evaluation of SCAD

#### Evaluation of immune stimulation in mouse CNS of ICV administrated SCAD-siRNA

To assess the potential immune stimulatory activity of SCAD-siRNA in the CNS, we measured the expression levels of two neuroinflammation markers—Iba1 (indicative of microglia activation) and Gfap (indicative of astrocyte activation) (Alterman et al., 2019, Belgrad et al., 2024)—in the hippocampus of mice. These mice received a single intracerebroventricular (ICV) dose of SCAD-si*Sod1* at 21 nmol, and samples were collected at various time points from day 14 to day 154 post-administration. As shown in Figure S3, there were no significant changes in the expression of either genes throughout the duration of the study, indicating that SCAD-siRNA does not elicit immune stimulation in the CNS.

#### Toxicity in rats following a single intrathecal injection

Safety and tolerability of the SCAD-siRNA were further assessed in Sprague-Dawley rats over a 14-day period following a single IT dose of two SCAD-conjugated siRNAs (SCAD-siRNA1 and SCAD-siRNA2) which did not have conserved target for any rat gene at doses of 1.6 mg or 4.8 mg. Throughout the observation period, there were no instances of body weight loss or any clinical signs indicative of distress or toxicity in any of the treatment groups compared to the control group treated with aCSF (Table S2). The comprehensive evaluation revealed no significant observational or statistical deviations across the treatment and control groups in terms of mortality, body weight, and clinical signs (Fig. S4).

Laboratory assessments including complete blood count (CBC), serum chemistry, and protein levels in the CSF showed no abnormalities (Fig. S5, S6, S7), confirming the absence of systemic toxicity. During functional observational battery (FOB) assessments, a slight decrease in neuromuscular function was noted in 1 out of 10 rats in the high-dose groups at day 1 and day 14, indicating potential test article related neurotoxicity (Fig. S8).

Histopathological examinations of the kidney revealed mild inflammation and dilation of the renal pelvis in a few animals, occurrences that were also noted in the control group and displayed no dose-dependent trends (Fig. S9). Notably, two animals receiving the lower dose of SCAD-siRNA1 exhibited mild degeneration of the spinal cord, likely related to the specific location of the intrathecal injection site or potentially due to procedural complications during CSF collection. These findings suggest that the observed histopathological changes were incidental and not related to SCAD-siRNA treatment.

#### Acute toxicity and immune stimulation of IV administrated SCAD-siRNA in ICR mice

To assess the acute toxicity and potential immune stimulation of IV administered SCAD-siRNA in ICR mice, we evaluated a duplex siRNA (si*SOD1)*, the ACO linked with the spacer 9 (ACO+S9), and a SCAD-siRNA (SCAD-si*SOD1*) at doses of 10.18 nmol and 40.71 nmol. We monitored serum chemistry, cytokine levels (IL-1β, IFN-γ, and TNFα), and mRNA expression for immune-related genes (*Ccl2*, *Ccl5*, *Lfit1*, *Lfit2*, *Xccl10*, *Ifih1*, *Isg15*) 8 hours after dosing. Clinically, one mouse in the low-dose siRNA group exhibited increased heart rate, paralysis, and mortality approximately one-hour post-injection, most likely procedure related. Serum levels of ALT, AST, and creatinine remained within normal ranges across all groups (Fig. S10A), although ALT levels were notably higher in the SCAD-si*SOD1* low-dose group compared to saline treatment. None of the treatments induced cytokine production (Fig. S10B). Regarding immune response genes, there was no significant activation in the spleen or liver across treatments compared to saline controls (Fig. S10C-D); however, Isg15 was significantly downregulated by SCAD-si*SOD1* at low dose in the spleen, and by both ACO+S9 at high dose and SCAD-si*SOD1* at low dose in the liver (Fig. S10C-D). The absence of dose-dependence suggests that this downregulation might be incidental, warranting further investigation.

## Discussion

Despite the FDA approval of six siRNA drugs targeting the liver, no duplex RNA drugs have yet received approval for CNS targets, primarily due to the formidable challenge of delivering these molecules to the CNS. The development of the SCAD system marks a substantial advancement in the delivery of duplex RNAs to the CNS and other extrahepatic tissues. The SCAD system utilizes the self-delivery properties of ASOs by incorporating an accessory oligonucleotide (ACO) to enhance the distribution and cellular uptake of duplex RNAs. Our results reveal that SCAD-siRNA facilitates profound and sustained knockdown across various CNS regions when administered locally, while also demonstrating a favorable safety profile. Additionally, we observed significant delivery efficacy outside of the CNS by local administration including the lung, the eye and the joint. Owing to its modular design, along with straightforward scalability and manufacturability, the SCAD architecture can be seamlessly adapted to any duplex RNA, as demonstrated with our mouse cell-type specific knocking down of *Map2*, *Gfap* and *Iba1*, thus expanding its potential to address a wide spectrum of diseases.

A critical factor in the success of SCAD is the optimization of its ACO component. Our research has shown that its sequence context, length and chemical modifications—particularly the PS backbone—play crucial roles in enhancing SCAD’s delivery efficiency and gene-silencing efficacy. Additionally, the choice of linkers connecting the ACO to the dsRNA is important, underscoring the necessity for meticulous component design to maximize therapeutic outcomes. Since the actual sequence can impact SCAD activity, this may suggest that the ACO acts as an aptamer to interreact and to bind with specific proteins.

The pharmacokinetic profile of SCAD-siRNA post-ICV injection demonstrates effective CNS retention and rapid systemic clearance. SCAD-si*Sod1* achieves peak plasma concentrations within an hour of administration, indicative of swift migration from CNS to systemic circulation, followed by rapid clearance within a day. This quick transition through systemic pathways minimizes peripheral exposure, enhancing safety. The comparative data between plasma and liver suggest an initial rapid drainage from the CNS into the systemic circulation, followed by liver absorption, with SCAD-si*Sod1* exhibiting a higher initial concentration than si*Sod1*.

In the CNS, SCAD-siRNA maintains higher concentrations compared to conventional duplex siRNA, with substantial knockdown effects lasting over five months. This sustained activity across various CNS regions—from cortical areas to deep brain structures—reveals a strong correlation between tissue concentration and pharmacodynamic activity, supporting the potential for extended dosing intervals.

Overall, these results confirm SCAD’s ability to deliver efficient and prolonged siRNA activity within the CNS, demonstrating its suitability for long-term therapeutic applications in neurodegenerative diseases.

The cell-type specificity of SCAD-siRNA in the CNS, as demonstrated in our study, underscores its potential for treating neurodegenerative diseases, where the ability to target a broad range of cell types is critical. Our results indicate that SCAD-siRNA not only efficiently targets neurons but also significantly impacts astrocytes and microglia. This capacity is vital as the neuronal microenvironment, which encompasses these cell types, plays a pivotal role in the progression of neurodegenerative diseases (Vahsen et al., 2021, D’Souza et al., 2023). In ALS, for instance, the pathogenesis involves not just neuronal dysfunction but also complex interactions within this microenvironment (Vahsen et al., 2021, Baird et al., 2024). The synthesis of mutant SOD1 in motor neurons initiates disease, while its expression in glial neighbors accelerates disease progression, highlighting the non-cell autonomous nature of ALS associated with *SOD1* mutations (Polymenidou and Cleveland, 2011, Boillée et al., 2006). In line with this, mutant *SOD1*-expressing astrocytes have been identified as viable therapeutic targets for slowing the progression of motor neuron degeneration in ALS (Yamanaka et al., 2008, Di Giorgio et al., 2007). By effectively reducing target mRNA levels in these glial cells, SCAD-siRNA can directly target upstream events of the pathogenesis process and potentially mitigate the toxic effects of protein aggregation, thus enabling disease-modification. Overall, the ability of SCAD to enable significant siRNA activity in a wide variety of tissues and cell types, both potently and in a durable manner, underscored its utility in the oligo drug development space.

Moreover, SCAD’s proficiency in delivering siRNAs to tissues beyond the CNS—such as the lung, the eye and the joint—broadens its application to other diseases where localized delivery of oligonucleotides is feasible.

The SCAD system’s ability to maintain high tissue concentrations and sustained gene knockdown over an extended period of time in the CNS highlights its potential for long-term therapeutic applications, with an infrequent dosing schedule. Additionally, the minimal systemic exposure and rapid clearance of SCAD-siRNA from peripheral tissues support its favorable safety profile.

SCAD demonstrated a favorable safety profile, showing no significant immune stimulation or systemic toxicity in our evaluations. In mice administered SCAD-si*Sod1* via ICV, neuroinflammation markers *Iba1* and *Gfap* remained stable, indicating no immune activation in the CNS. Similarly, in Sprague-Dawley rats, a single intrathecal injection of SCAD-siRNA resulted in no adverse clinical signs, body weight loss, or significant deviations in laboratory tests, including CBC, serum chemistry, and CSF protein levels, when compared to controls. Minor decreases in neuromuscular function were observed at the highest dose but occurred at a very low frequency (1 out of 10 rats), suggesting a possible, albeit minimal, impact of the treatment.

Histopathological analysis revealed mild renal changes and isolated spinal cord degeneration, both of which were non-dose-dependent. The spinal cord degeneration was considered likely related to the procedure (IT injection), whereas the renal changes appeared incidental and not directly related to the test article. In ICR mice, intravenous administration of SCAD-siRNA showed no significant cytokine induction or immune gene activation, though a minor, incidental increase in serum ALT levels was noted in one group. Overall, these findings reinforce the safety of SCAD, supporting its potential for further clinical development in CNS and other localized therapeutic applications.

In conclusion, the SCAD delivery platform emerges as a promising solution for the efficient delivery of oligonucleotide therapeutics to challenging target tissues. Its capacity to enhance distribution, cellular uptake, and sustained gene knockdown positions SCAD as an invaluable tool in developing novel treatments for genetic disorders, including neurodegenerative diseases. The ongoing clinical trial for *SOD1*-ALS will provide essential insights into the safety and efficacy of SCAD-enabled siRNA therapy, setting the stage for future clinical applications.

## Material and methods

### Oligonucleotides synthesis

Oligonucleotides for both *in vitro* and *in vivo* studies were synthesized in-house as listed in Supplementary Table 1. Single-stranded oligonucleotides were synthesized on a solid-phase support using an HJ-12 synthesizer (Highgene-Tech Automation, Beijing, China) and purified by reverse-phase high-performance liquid chromatography (RP-HPLC) with an acetonitrile gradient over a UniPS column (NanoMicro Technology, Suzhou, China). Each sequence was then reconstituted in sterile water. Equal amounts of each strand were annealed into duplexes by heating and then cooling to room temperature. Resolution of a single band via gel electrophoresis at the predicted molecular weights was used to evaluate duplex formation. Electrospray ionization mass spectrometry (ESI-MS) confirmed the duplex identity, while overall purity was analyzed via size-exclusion chromatography high-performance liquid chromatography (SEC-HPLC) using an XBridge Protein BEH SEC 125 A column (Waters Corporation, Milford, MA, USA). Endotoxin levels in each batch were quantified using the endpoint Chromogenic Endotoxin Quant Kit (Bioendo, Xiamen, China) via proenzyme Factor C.

### Electrophoretic mobility shift assays (EMSA)

Oligonucleotides were incubated at 37°C in 50% mouse plasma at a final concentration of 0.5 μM for 1 hour. Samples were mixed with loading buffer and loaded onto a 4% agarose gel. Electrophoresis was performed at 120 volts for 60 minutes. After electrophoresis, the gel was documented using a ChemiDoc MP system to quantify the intensity of each oligonucleotide band. The oligonucleotide band from the plasma-mixed oligonucleotide sample with the same migration rate on the gel as their input was regarded as the unbound fraction. For each sample, a percentage of the unbound fraction was calculated as follows: Unbound fraction (%) = (Intensity of oligonucleotide band from plasma-mixed sample) / (Intensity of oligonucleotide band from input) × 100.

### Primary mouse hepatocyte (PMH) isolation and oligonucleotide treatment

PMHs were isolated from C57BL/6J mice using a modified method (Charni-Natan and Goldstein, 2020). Mice were anesthetized and perfused with initial flushing reagent and digestion reagent. The liver was then dissected and mechanically dissociated. The cell suspension was filtered and centrifuged. Cells with at least 80% viability were seeded onto collagen I-coated plates and allowed to reach 90-95% confluence before the assay. PMH cells were cultured under standard conditions in modified William’s Medium E (Gibco) supplemented with insulin and penicillin/streptomycin. Mock (control) and pre-mixed oligonucleotides with 50% mouse serum were added to the medium at 100 or 1000 nM for three days without transfection reagents to facilitate passive uptake, followed by Lipofectamine RNAiMAX transfection (Thermo Fisher) following the manufacturer’s instructions.

### Animal procurement

All animal procedures followed local and state regulations and were approved by the Institutional Animal Care and Use Committee of Ractigen Therapeutics. C57BL/6 mice (4-6 weeks old) were obtained from JOINN Biologics (Jiangsu, China) and Sprague-Dawley (SD) rats (4-6 weeks old) were purchased from Nantong University (Jiangsu, China).

### Gene expression analysis via RT-qPCR

Animal tissues were homogenized in RNAVzol (Biosharp, China). RNA was isolated using the MagPure Total RNA Micro LQ kit (Magen, China). RNA concentrations were determined, and reverse transcription was performed using the PrimeScript RT kit (Takara) with gDNA Eraser. cDNA was amplified by real-time PCR using SYBR Premix Ex Taq II (Takara) with specific primers. Melting curves confirmed primer specificity. Relative gene expression was calculated using the ΔΔCt method. Primer sequences are available in Supplementary Table 2.

### Quantification of siRNA in animal tissues

Quantification of siRNA in animal tissues involves preparing tissue lysate by a Bioprep-24 Homogenizer (Allsheng). To inactivate sample proteins, the lysates are heated to 95°C. An 8-point standard curve is generated using serial dilutions of non-treated lysate spiked with si*Sod1* or SCAD. Reverse transcription (RT) reactions are subsequently performed custom stem-loop primers and amplified specific to siRNA guide strands by real-time PCR (Supplementary Table 3). Finally, absolute quantities of siRNA are extrapolated by linear regression using the standard curves. Tissue concentrations are then calculated as the ratio of absolute siRNA mass (ng) to the total weight (g) of the tissue sample used for lysis preparation.

### Intracerebroventricular administration

Mice were anesthetized with Avertin and placed in a stereotaxic apparatus. A scalp incision was made, and a needle containing the siRNA formulation was positioned at the bregma. The needle was inserted into the lateral ventricle, and 10 μL of solution was injected at a rate of 1 μL/s. The wound was then sutured closed.

### Intrathecal administration

SD rats were anesthetized with isoflurane. The injection site at the base of the tail was shaved and cleaned. A needle was carefully inserted into the intradural space between the L5-L6 spinous processes until a tail flick was observed. A total volume of 10 μL of solution was injected over 1 minute.

### Intratracheal administration

For intratracheal administration, modified from a previous method (Ortiz-Munoz and Looney, 2015), anesthetized mice were placed in a supine position. A small incision was made at the neck to expose the trachea. Oligonucleotides (no more than 50 µL) were slowly administered into the lung. The incision was then closed with sutures.

### Intravitreal administration

SD rats were anesthetized with isoflurane. An anterior chamber paracentesis allowed some aqueous humor to drain. The oligonucleotides, dissolved in 4 µL of normal saline, were loaded into a 30-gauge needle for IVT injection at a 45° angle into the posterior chamber. After administration, antibiotics were applied to the injected eye to prevent infection.

### Intraarticular administration

The intra-articular administration process for rat knee joints, modified from a previous method (Adaes et al., 2014). After anesthesia and sterilizing the knee area, a needle was inserted into the joint. A standard volume of 30 μL was administered, followed by gentle flexion and extension of the knee joint to aid in the distribution of the injected material. Post-injection, the rats were monitored for any signs of distress or complications.

### Clinical observation and endpoint criteria

Cage-side clinical signs, i.e., poor health, behavioral changes et.al. were recorded at least once a day during the acclimation period and twice a day (AM and PM) during the treatment. Abnormal symptoms (including convulsions, seizures, rigidity, muscle tremor, and dyspnea), if occur, were described in details and videos or photos were taken. Animals were observed for 4 h after injection and daily thereafter until endpoint. Body weight was recorded before test substance administration and at recorded intervals thereafter. Humane endpoint criteria were defined as weight loss >20% relative to their initial mass at day of treatment or when animals reach Neuroscores bigger or equals to 4.

### Statistical analysis

Data analysis was performed using GraphPad Prism version 10.1.2 (324). Differences in continuous variables between treatments were assessed by a two-tailed and unpaired Student’s *t*-test (for two treatments) or *One-way* ANOVA followed by Dunnett’s test for multiple comparison corrections (for three or more treatments). Time-stratified data (such as peak weight analysis and animal survival) were plotted using Kaplan-Meier graphs. Significance was defined as follows: *, *P* < 0.05; **, *P* < 0.01; ***, *P* < 0.001; ****, *P* < 0.0001.

