## Supplementary materials for "SCAD Delivery Platform: A Novel Approach for Efficient CNS and Extrahepatic Oligonucleotide Therapeutics"

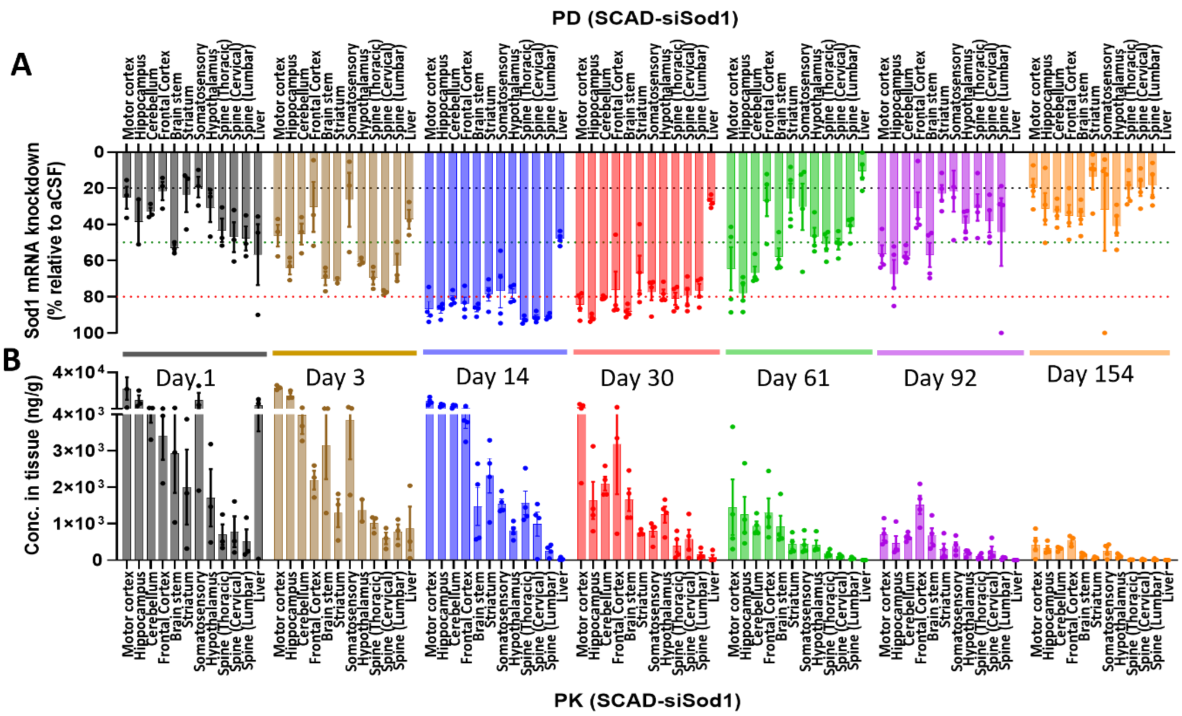


**Supplementary Figure 1. SCAD-siRNA achieved broad tissue exposure and durable knockdown activity in the CNS of C57BL/6J mice**. The PD (**A**) and PK (**B**) data from Figure 5 is plotted as bar graphs for better visualization. The red, green and grey dotted line in (**A**) denote 80%, 50% and 20% knock-down respectively.

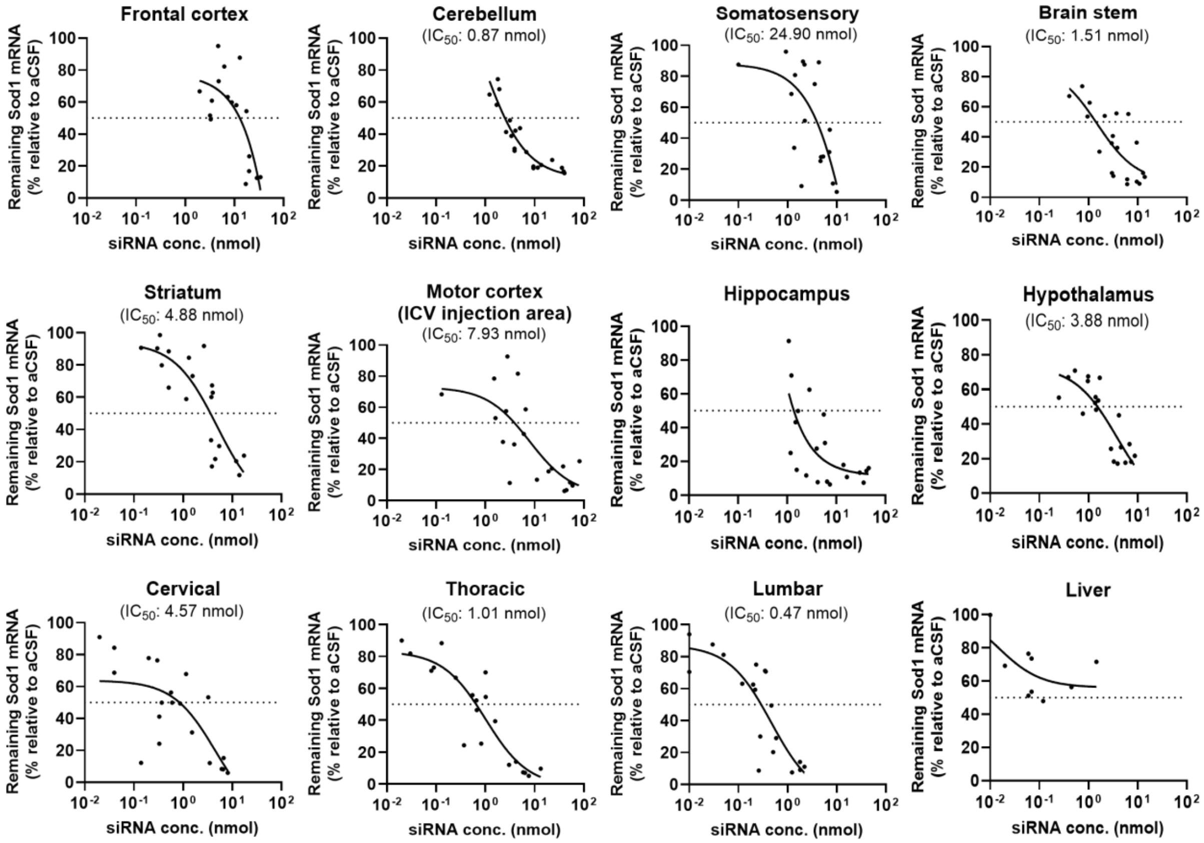


**Supplementary Figure 2. PD-PK relationship of SCAD-siRNA in C57BL/6J mice**. The mice were administrated with SCAD-si*Sod1* at 21 nmol or received aCSF as a vehicle control via ICV injection. The concentration of siRNA guide strand and the expression of *Sod1* were analyzed in various tissues in the brain (*i.e.*, frontal cortex, cerebellum, somatosensory, brain stem, striatum, motor cortex, hippocampus, and hypothalamus), as well as the spinal cord (*i.e.*, cervical, thoracic, and lumbar), and liver were analyzed at different time points (day 14, 30, 61, 92 and 154) after dosing using stem-loop RT-qPCR and RT-qPCR, respectively.

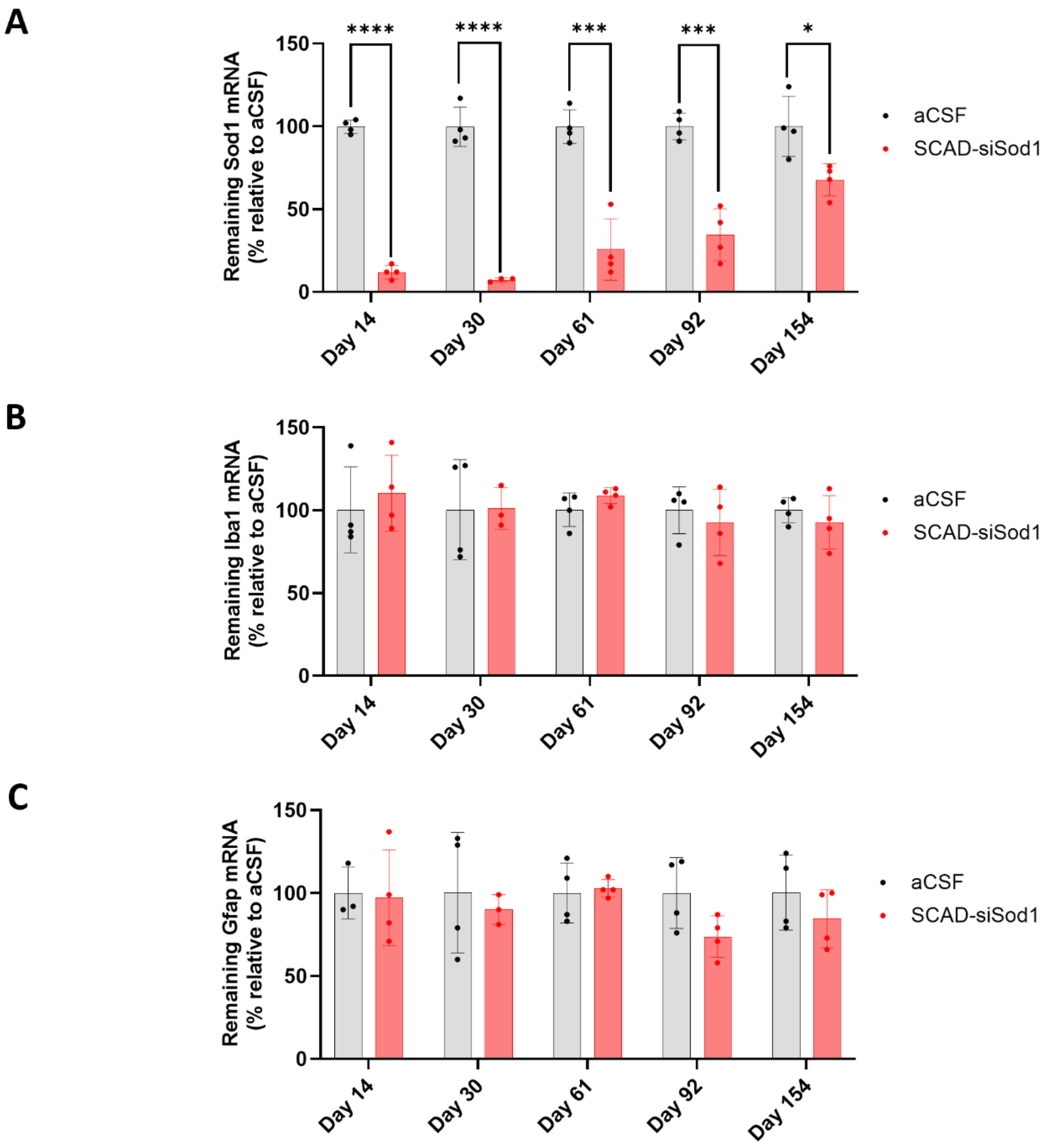


**Supplementary Figure 3. SCAD-siRNA did not induce neuroinflammation in mouse hippocampus.** C57BL/6J mice were administrated with SCAD-siSod1 at a molar dose of 21 nmol via ICV injection or received aCSF as a vehicle control. The expression of *Sod1* (**A**), *Iba1* (**B**), and *Gfap* (**C**) were analyzed in hippocampus tissues at different time points over a 154-day period using RT-qPCR. The data represent the mean values ± SD, with n=3-4 mice per group.

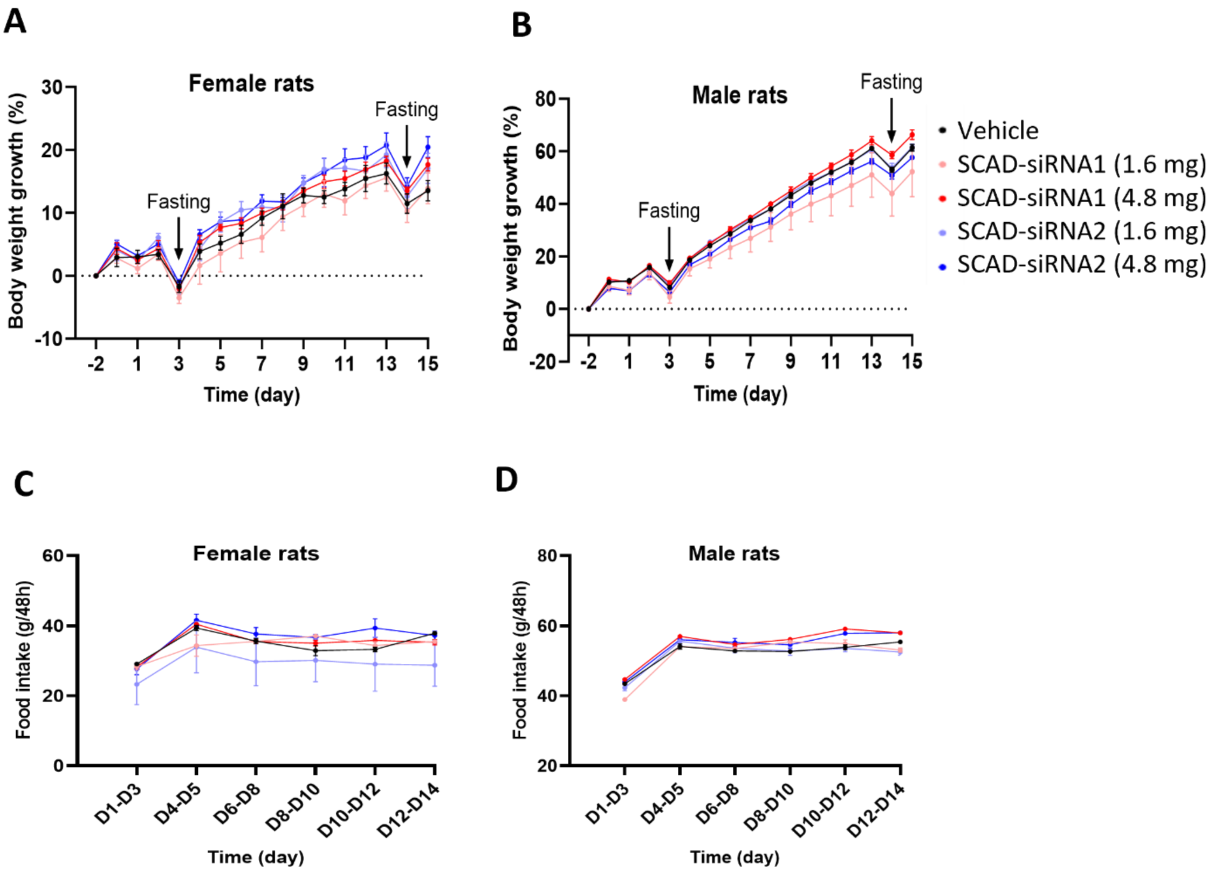


**Supplementary Figure 4. Body weight change and food intake following IT dosing of SCAD-siRNA in rats.** Rats received either SCAD-siRNA1 or SCAD-siRNA2 at doses of 1.6 mg or 4.8 mg via IT injection. Each treatment group comprised 10 rats, evenly divided between 5 males and 5 females, with a vehicle control group receiving aCSF. Fasting was carried out on day 3 and 14 prior to the collection of serum for serum biochemistry assay. **A** and **B**. Body weight change. **C** and **D**. Food intake measured in grams.

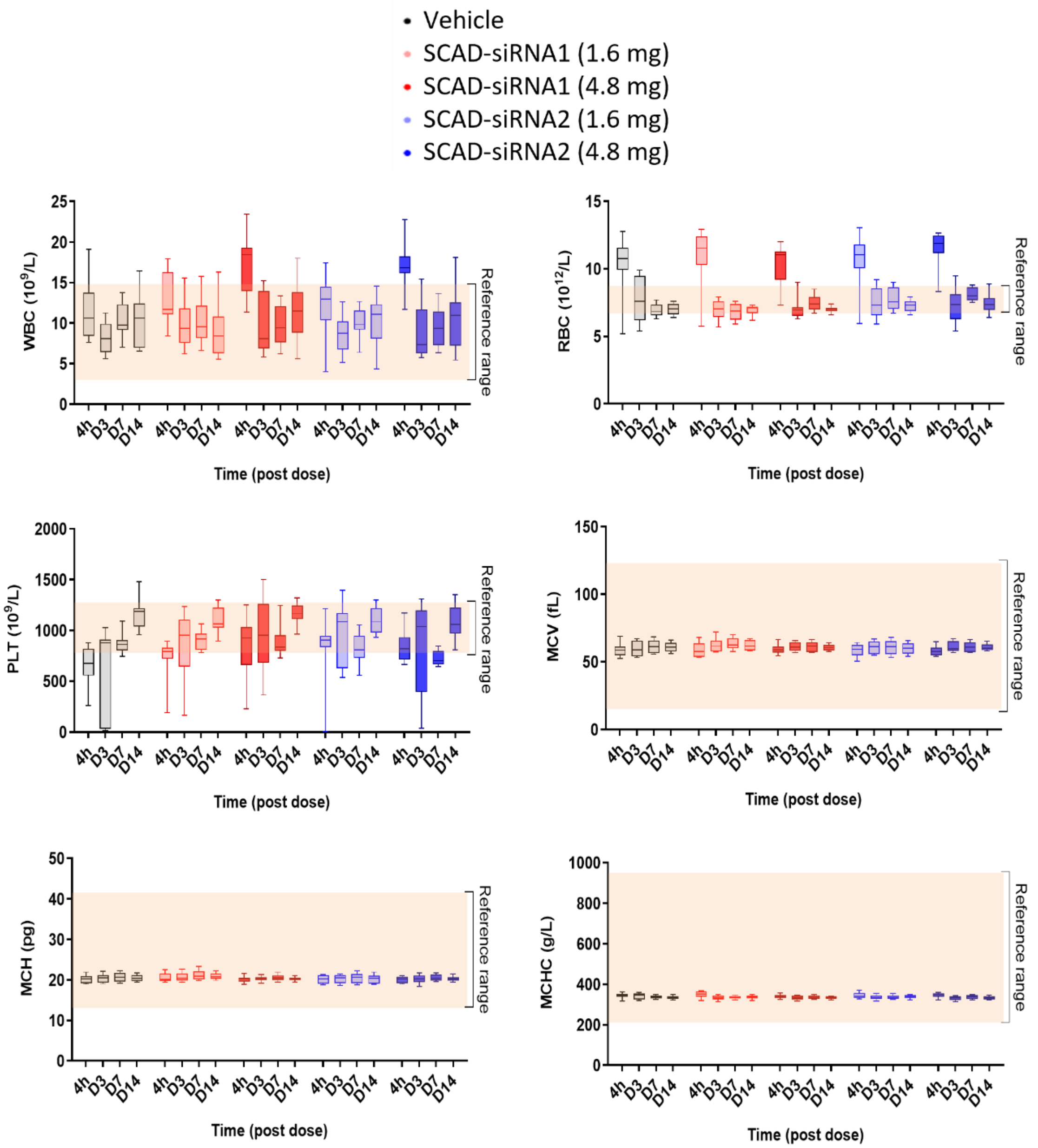


**Supplementary Figure 5. Complete blood count (CBC) analysis post IT-administration of SCAD-siRNA in Rats.** Rats received either SCAD-siRNA1 or SCAD-siRNA2 at doses of 1.6 mg or 4.8 mg via IT injection. Each treatment group comprised 10 rats, evenly divided between 5 males and 5 females, with a vehicle control group receiving aCSF. CBC analysis was performed on non-anticoagulated whole blood collected at 4 hours, 3 days, 7 days, and 14 days after dosing. Parameters analyzed included white blood cells (WBC), red blood cells (RBC), platelets (PLT), mean cell volume (MCV), mean cell hemoglobin (MCH), and mean cell hemoglobin concentration (MCHC).

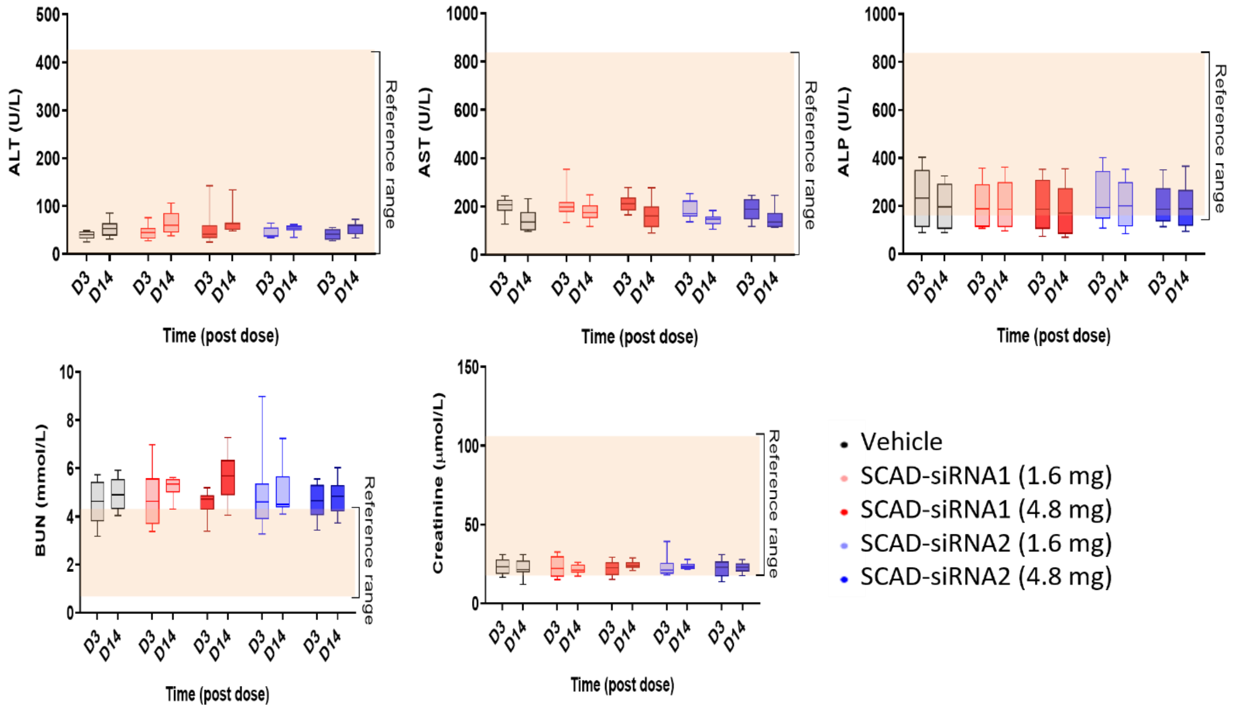


**Supplementary Figure 6. Serum chemistry analysis conducted (IT) injection of SCAD in rats.** Rats received either SCAD-siRNA1 or SCAD-siRNA2 at doses of 1.6 mg or 4.8 mg via IT injection. Each treatment group comprised 10 rats, evenly divided between 5 males and 5 females, with a vehicle control group receiving aCSF. Blood samples were collected at two time points (3 and 14 days post-administration) to assess ALT, AST, and ALP as indicators of liver function, and BUN and creatinine as markers of kidney function.

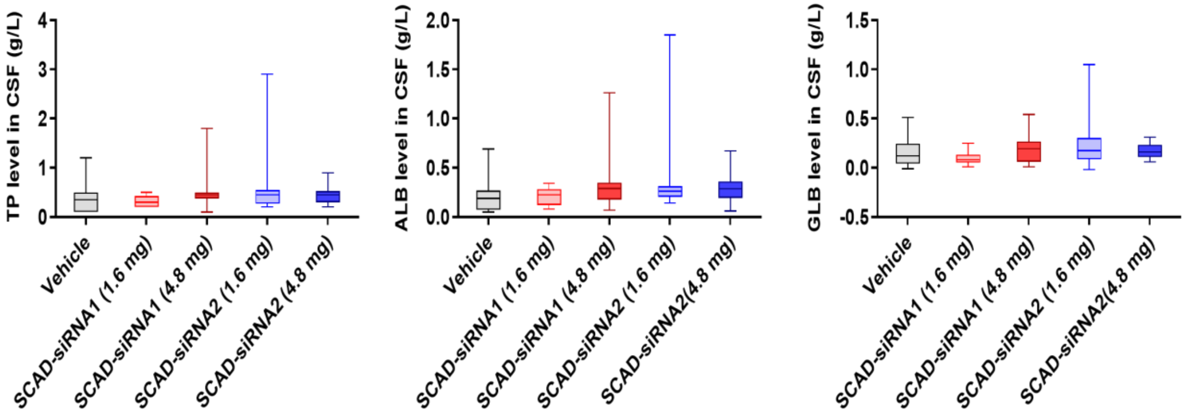


**Supplementary Figure 7. CSF protein levels conducted 14 days posting IT dosing of SCAD-siRNA in rats.** Rats received either SCAD-siRNA1 or SCAD-siRNA2 at doses of 1.6 mg or 4.8 mg via IT injection. Each treatment group comprised 10 rats, evenly divided between 5 males and 5 females, with a vehicle control group receiving aCSF. CSF was collected from the animals after 14 days, and total protein (TP), albumin (ALB), and globulin (GLB) levels were determined using a biochemical analyzer (Olympus AU640 Chemistry Analyzer).

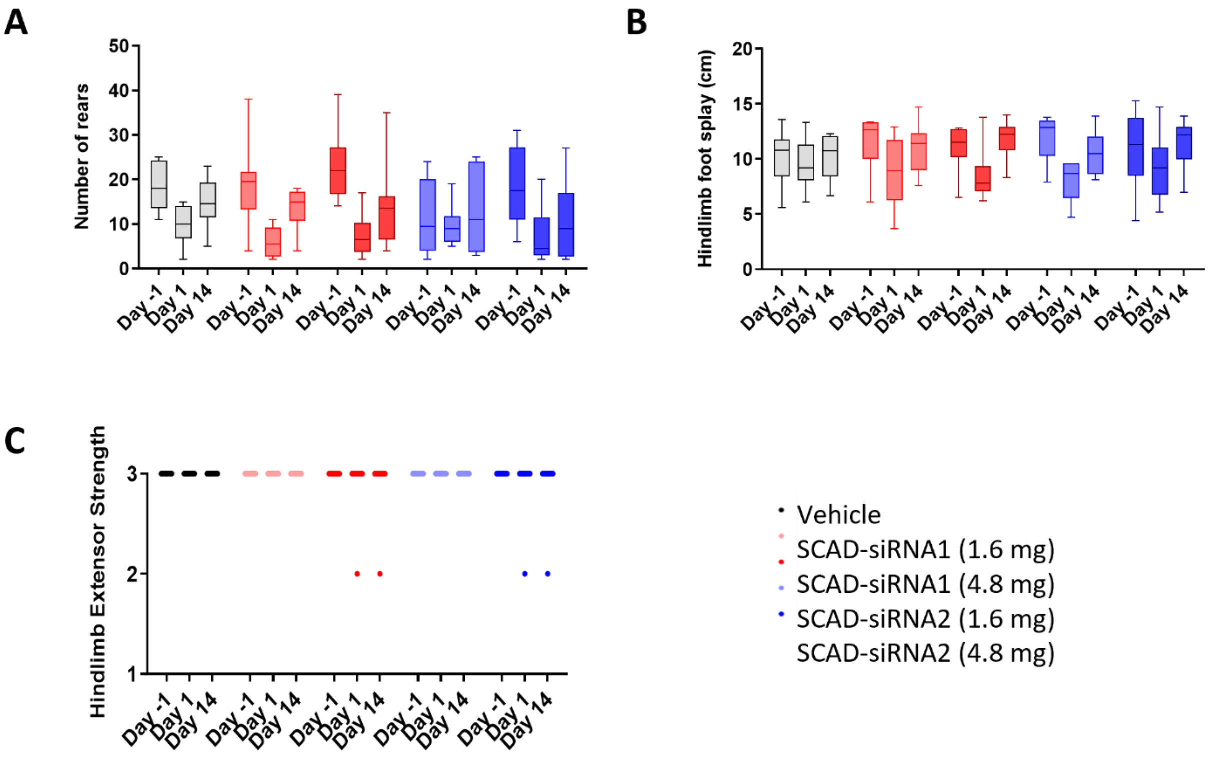


**Supplementary Figure 8. Functional observational battery (FOB) scores in rats before and after receiving an IT dosing of SCAD-siRNA.** Rats received either SCAD-siRNA1 or SCAD-siRNA2 at doses of 1.6 mg or 4.8 mg via IT injection. Each treatment group comprised 10 rats, evenly divided between 5 males and 5 females, with a vehicle control group receiving aCSF. FOB was assessed on Day -1 (pre-dosing), Day 1 (3 to 8 hours post-dosing), and Day 14 (endpoint). The assessment included: **A**. the number of rear-ups on hind legs, **B**. hindlimb foot splay, and **C**. hindlimb extensor strength, which was rated as 1 for absent, 2 for reduced, or 3 for normal.

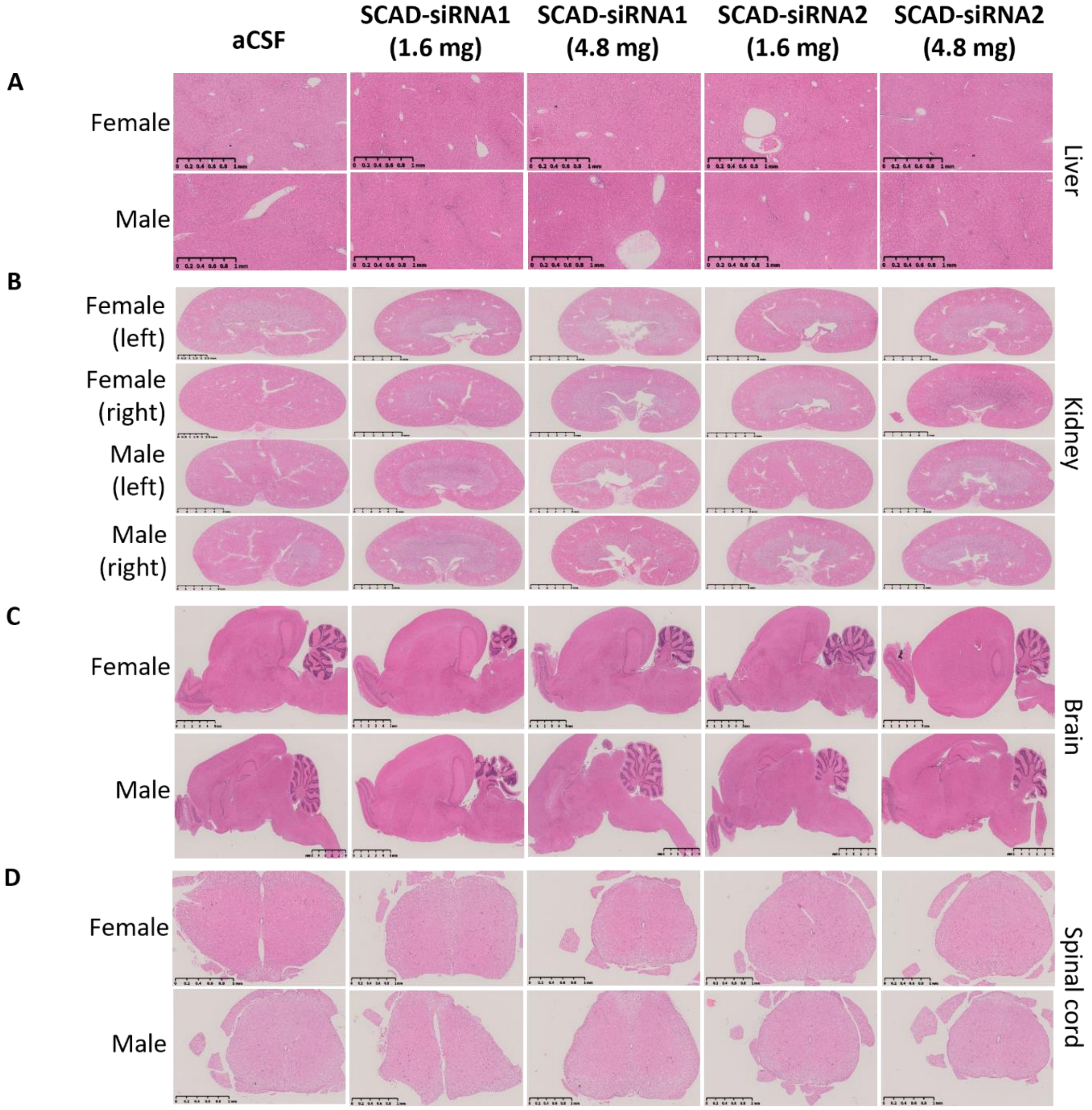


**Supplementary Figure 9. The results of a histopathological examination conducted after the intrathecal (IT) injection of SCAD in rats.** SD rats received either SCAD-siRNA1 or SCAD-siRNA2 at doses of 1.6 and 4.8 mg. Each group consisted of 10 rats, with an equal distribution of 5 males and 5 females. The injections were performed intrathecally. The vehicle control group was administrated aCSF. Organs/tissues were collected and stained with hematoxylin and eosin (H&E). This figure presents representative histology images of liver (**A**), kidney (**B**), brain (**C**), and spinal cord (**D**).

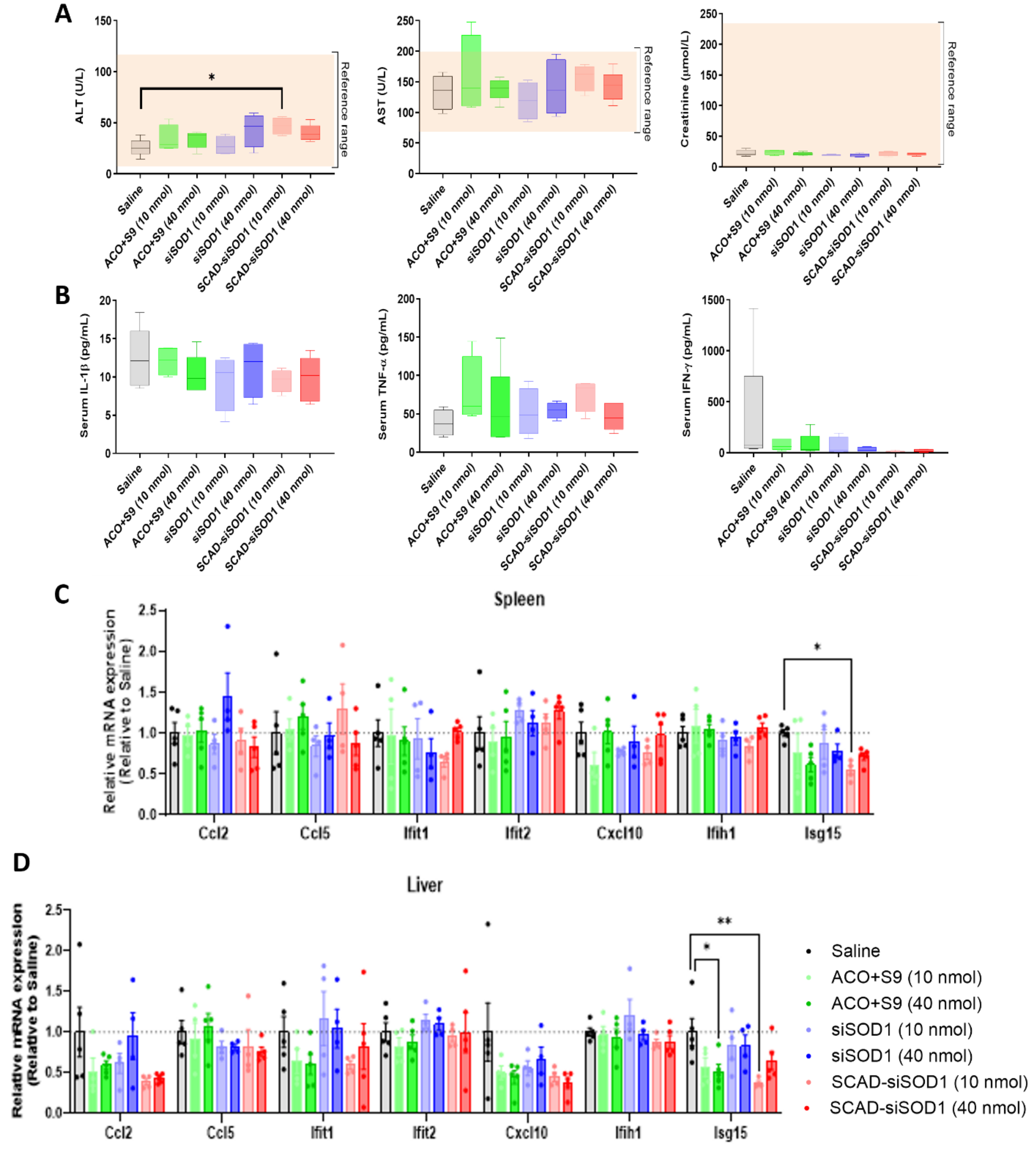


**Supplementary Figure 10. Acute systemic toxicity via intravenous (IV) injection of SCAD and SCAD components in mice.** ICR mice received ACO+S9 alone, si*SOD1* alone, and SCAD-si*SOD1* at doses of 10 or 40 nmol via IV injection. Each treatment group comprised 4-5 female mice, with a vehicle control group receiving saline and being sacrificed 8 hours post-dose. **A**. Serum samples were assessed for ALT and AST as indicators of liver function, and creatinine as markers of kidney function. **B**. Immune responses were analyzed for IL-1b, TNF-a, IFN-g using ELISA kits. **C** and **D**. Potential immune response gene expression was analyzed in spleen (C) and liver(D) tissues by RT-qPCR using *Ccl2*, *Ccl5*, *Ifit1*, *Ifit2*, *Cxcl10*, *Ifih1*, and *Isg15* specific primers.

**Supplementary table 1. Single ICV dose PK study.**

| **Tissue** | **Concentration (ng/µg, means ± SEM)** | | | | | | | | | |
|---|---|---|---|---|---|---|---|---|---|---|
| | | **1 h** | **8 h** | **D1** | **D3** | **D14** | **D30** | **D61** | **D92** | **D154** |
| Frontal cortex | siSod1 | 11724 ±4946 | 4768 ± 2419 | 4856 ± 3824 | 992 ± 275 | 3423 ± 914 | 1281 ± 498 | 515 ± 229 | 305 ± 92 | 10 ± 3 |
| | SCAD-siSod1 | 2608 ± 1466 | 5120 ± 1365 | 3415 ± 656 | 2195 ± 256 | 4341 ± 729 | 3180 ± 1371 | 1295 ± 402 | 1516 ± 253 | 548 ± 89 |
| | **Ratio*** | **0.22** | **1.07** | **0.70** | **2.21** | **1.27** | **2.48** | **2.51** | **4.97** | **54.80** |
| Motor cortex | siSod1 | 56783 ± 29782 | 44074 ± 16603 | 5254 ± 1579 | 5110 ± 127 | 8047 ± 757 | 3120 ± 1624 | 646 ± 330 | 260 ± 78 | 86 ± 55 |
| | SCAD-siSod1 | 119996 ± 43644 | 49484 ± 31477 | 24096 ± 11147 | 24891 ± 1298 | 11130 ± 1661 | 5504 ± 1427 | 1455 ± 762 | 705 ± 162 | 428 ± 180 |
| | **Ratio** | **2.11** | **1.12** | **4.59** | **4.87** | **1.38** | **1.76** | **2.25** | **2.71** | **4.98** |
| Somatosen-sory cortex | siSod1 | 47706 ± 25156 | 12296 ± 5341 | 5342 ± 2288 | 1147 ± 166 | 1541 ± 360 | 587 ± 300 | 148 ± 56 | 123 ± 107 | 7 ± 2 |
| | SCAD-siSod1 | 11701 ± 6458 | 24258 ± 20070 | 12138 ± 7529 | 3854 ± 1085 | 1553 ± 127 | 793 ± 145 | 432 ± 136 | 376 ± 117 | 254 ± 96 |
| | **Ratio** | **0.25** | **1.97** | **2.27** | **3.36** | **1.01** | **1.35** | **2.92** | **3.06** | **36.29** |

| | | | | | | | | | | |
|---|---|---|---|---|---|---|---|---|---|---|
| Hypo-thalamus | siSod1 | 23510 ± 13596 | 3282 ± 1548 | 514 ± 266 | 624 ± 279 | 365 ± 110 | 188 ± 58 | 48 ± 20 | 10 ± 1 | 8 ± 2 |
| | SCAD-siSod1 | 22837 ± 2723 | 5482 ± 3095 | 1715 ± 789 | 1365 ± 302 | 810 ± 111 | 1249 ± 222 | 421 ± 124 | 197 ± 45 | 119 ± 30 |
| | **Ratio** | **0.97** | **1.67** | **3.34** | **2.19** | **2.22** | **6.64** | **8.77** | **19.70** | **14.88** |
| Brain stem | siSod1 | 35791 ± 13103 | 6728 ± 1998 | 3413 ± 1229 | 1821 ± 347 | 1336 ± 484 | 470 ± 62 | 394 ± 233 | 122 ± 26 | 92 ± 33 |
| | SCAD-siSod1 | 9920 ± 2406 | 5515 ± 3661 | 2938 ± 1093 | 3154 ± 931 | 1478 ± 525 | 1657 ± 311 | 921 ± 301 | 672 ± 206 | 151 ± 27 |
| | **Ratio** | **0.28** | **0.82** | **0.86** | **1.73** | **1.11** | **3.53** | **2.34** | **5.51** | **1.64** |
| Striatum | siSod1 | 13871 ± 6025 | 2803 ± 312 | 772 ± 79 | 399 ± 165 | 1179 ± 95 | 518 ± 333 | 43 ± 27 | 51 ± 9 | 8 ± 3 |
| | SCAD-siSod1 | 849 ± 416 | 1264 ± 411 | 2010 ± 1029 | 1295 ± 387 | 2320 ± 476 | 751 ± 35 | 435 ± 127 | 301 ± 155 | 61 ± 14 |
| | **Ratio** | **0.06** | **0.45** | **2.60** | **3.25** | **1.97** | **1.45** | **10.12** | **5.90** | **7.63** |
| Hippo-campus | siSod1 | 140082 ± 66138 | 23406 ± 7915 | 7039 ± 359 | 3035 ± 1292 | 3101 ± 514 | 603 ± 289 | 180 ± 77 | 141 ± 45 | 111 ± 57 |
| | SCAD-siSod1 | 84063 ± 15561 | 40006 ± 28336 | 12506 ± 4767 | 17131 ± 2408 | 7265 ± 633 | 1642 ± 513 | 1249 ± 490 | 465 ± 199 | 322 ± 75 |
| | **Ratio** | **0.60** | **1.71** | **1.78** | **5.64** | **2.34** | **2.72** | **6.94** | **3.30** | **2.90** |
| Cerebellum | siSod1 | 40432 ± 18710 | 4083 ± 1058 | 1915 ± 123 | 1641 ± 56 | 5862 ± 915 | 1604 ± 221 | 975 ± 267 | 286 ± 30 | 81 ± 26 |
| | SCAD-siSod1 | 50984 ± 15520 | 4207 ± 1985 | 4244 ± 469 | 3986 ± 521 | 6573 ± 761 | 2102 ± 188 | 935 ± 128 | 630 ± 57 | 315 ± 29 |
| | **Ratio** | **1.26** | **1.03** | **2.22** | **2.43** | **1.12** | **1.31** | **0.96** | **2.20** | **3.89** |

| | | | | | | | | | | |
|---|---|---|---|---|---|---|---|---|---|---|
| Spinal cord (cervical) | siSod1 | 7640 ± 2618 | 731 ± 397 | 631 ± 517 | 254 ± 68 | 365 ± 42 | 185 ± 43 | 110 ± 40 | 30 ± 4 | 1 ± 1 |
| | SCAD - siSod1 | 1230 ± 452 | 364 ± 255 | 781 ± 425 | 597 ± 165 | 991 ± 332 | 571 ± 264 | 102 ± 23 | 251 ± 126 | 14 ± 8 |
| | **Ratio** | **0.16** | **0.50** | **1.24** | **2.35** | **2.72** | **3.09** | **0.93** | **8.37** | **14.00** |
| Spinal cord (thoracic) | siSod1 | 8572 ± 4161 | 859 ± 335 | 676 ± 487 | 361 ± 111 | 444 ± 114 | 172 ± 134 | 22 ± 6 | 15 ± 5 | 2 ± 2 |
| | SCAD - siSod1 | 1544 ± 289 | 360 ± 176 | 701 ± 282 | 1022 ± 140 | 1574 ± 325 | 401 ± 173 | 181 ± 44 | 98 ± 38 | 10 ± 3 |
| | **Ratio** | **0.18** | **0.42** | **1.04** | **2.83** | **3.55** | **2.33** | **8.23** | **6.53** | **5.00** |
| Spinal cord (lumbar) | siSod1 | 8203 ± 4067 | 827 ± 353 | 842 ± 539 | 387 ± 102 | 120 ± 26 | 57 ± 32 | 26 ± 14 | 12 ± 7 | 1 ± 0 |
| | SCAD - siSod1 | 895 ± 144 | 207 ± 115 | 523 ± 324 | 782 ± 203 | 262 ± 81 | 151 ± 65 | 48 ± 15 | 56 ± 16 | 14 ± 10 |
| | **Ratio** | **0.11** | **0.25** | **0.62** | **2.02** | **2.18** | **2.65** | **1.85** | **4.67** | **14.00** |
| Liver | siSod1 | 158 ± 30 | 486 ± 436 | 169 ± 149 | 207 ± 191 | 0 ± 0 | 0 ± 0 | 0 ± 0 | 0 ± 0 | 0 ± 0 |
| | SCAD - siSod1 | 56 ± 28 | 2843 ± 557 | 7480 ± 3944 | 870 ± 602 | 33 ± 18 | 77 ± 67 | 1 ± 0 | 0 ± 0 | 0 ± 0 |
| | **Ratio** | **0.35** | **5.85** | **44.26** | **4.20** | **n/a** | **n/a** | **n/a** | **n/a** | **n/a** |
| Kidney | siSod1 | 7300 ± 258 | 14988 ± 2614 | 9316 ± 2437 | 4574 ± 859 | | | | | |
| | SCAD - siSod1 | 4031 ± 930 | 38778 ± 4726 | 45854 ± 6920 | 8856 ± 3914 | | | | | |
| | **Ratio** | **0.55** | **2.59** | **4.92** | **1.94** | | | | | |
| Plasma | siSod1 | 17019 ± 2772 | 152 ± 96 | 1 ± 0 | 1 ± 0 | 0 ± 0 | 2 ± 1 | 0 ± 0 | 0 ± 0 | 0 ± 0 |

| | | | | | | | | | | |
|---|---|---|---|---|---|---|---|---|---|---|
| | SCAD - siSod1 | 69320 ± 14720 | 69 ± 12 | 3 ± 1 | 1 ± 0 | 6 ± 2 | 36 ± 12 | 0 ± 0 | 0 ± 0 | 0 ± 0 |
| | **Ratio** | **4.07** | **0.45** | **3.00** | **1.00** | **n/a** | **18.00** | **n/a** | **n/a** | **n/a** |
| * Ratio of SCAD-siSod1 to siSod1 | | | | | | | | | | |

**Supplementary table 2. Toxicology study of SCAD-siRNA in SD rats following a single IT dose with a 16-day observation period.**

| **Treatment** | **aCSF** | **SCAD$^{RD-12293}$** | **SCAD$^{RD-12293}$** | **SCAD$^{RD-12500}$** | **SCAD$^{RD-12500}$** |
|---|---|---|---|---|---|
| Dose | - | 1.6 mg | 4.8 mg | 1.6 mg | 4.8 mg |
| **Animal number** | 10 | 10 | 10 | 10 | 10 |
| **Mortality** | N/O | N/O | N/O | N/O | N/O |
| **Body weight loss** | N/O | N/O | N/O | N/O | N/O |
| **Clinical signs** | N/O | N/O | N/O | N/O | N/O |
| **CBC** | N/O | N/O | N/O | N/O | N/O |
| **Serum chemistry** | N/O | N/O | N/O | N/O | N/O |
| **Protein levels in CSF** | N/D | N/D | N/D | N/D | N/D |
| **FOB** | | | | | |
| Neuromuscular | N/O | N/O | Low hindlimb extensor strength (1/10) | N/O | Low hindlimb extensor strength (1/10) |
| Other | N/O | N/O | N/O | N/O | N/O |
| **Histopathology** | | | | | |
| Brain | N/C | N/C | N/C | N/C | N/C |
| Spinal cord | N/C | Steatosis (2/10) | N/C | N/C | N/C |
| Liver | N/C | N/C | N/C | N/C | N/C |
| Kidney | Renal pelvis (1/10) | N/C | Renal pelvis (3/10), Inflammation (1/10) | Renal pelvis (2/10), Inflammation (1/10) | N/C |
| Key: N/O: No observation; N/D: No statistical difference; N/C: No obvious pathological changes | | | | | |

**Supplementary table 3. Primer sequences.**

| **Primers** | **Forward (5'-3')** | **Reverse (5'-3')** |
|---|---|---|
| *SOD1* (human) | aagcattaaaggactgactgaagg | caagtctccaacatgcctctc |
| *Sod1* (mouse) | ttggccgtacaatggtggtc | acagtttaatggtttgagggtagc |
| *Sod1* (rat) | agcggatgaagagaggcatgt | cttcatttccacctttgccca |
| *Gfap* (mouse) | tcgcactcaatacgaggcag | ttggcggcgatagtcgttag |
| *Tubb3* (mouse) | agtgcggcaaccagatag | tccaagtccaccagaatg |
| *Iba1* (mouse) | tctgccgtccaaacttgaagcc | ctcttcagctctaggtgggtct |
| *Map2* (rat) | gggtggacactcaagatgatg | gagaccgtgctaggttcctt |
| *TBP* (human) | tgctcacccaccaacaatttag | tctgctctgactttagcacctg |
| *Tbp* (mouse) | ccgtgaatcttggctgtaaact | tgtccgtggctctcttattc |
| *RPL13a* (mouse) | caccctatgacaagaaaaagcg | tcttggccttttccttccgtt |
| *Tbp* (rat) | cttcgtgccagaaatgctga | tctggattgttcttcactcttgg |
| *Ccl2* (Mouse) | CTGGAGCATCCACGTGTTGG | TGAGCTTGGTGACAAAAACTACAGC |
| *Ifit1* (Mouse) | AGGAGTTCTGCTCTGCTGAAA | TCCCCAATGGGTTCTTGATGT |
| *Ifit2* (Mouse) | AGACTTAGAGGTGCTGCACAGA | GTCGCAGATTGCTCTCCAGT |
| *Cxcl10* (Mouse) | CACCATGAACCCAAGTGCTG | TCTTTTTCATCGTGGCAATGATCT |
| *Ifih1* (Mouse) | GACACCAGAATTCAAGGGACTCT | GTGCACCATCATTGTTCCCC |
| *Isg15* (Mouse) | AGCAGACTCCTTAATTCCAGGG | CCGTCATGGAGTTAGTCACG |
| *Ccl5* (Mouse) | TGCCCACGTCAAGGAGTATT | ACTTGGCGGTTCCTTCGAG |
| **Stem-loop RT-qPCR Primers (5'-3')** | | |
| Stem-loop RT | gtcgtatccagtgcagggtccgaggtattcgcactggatacgactcattt | |
| Stem-loop qPCR | cgcgcttagagtgaggattaaaa | gtgcagggtccgaggt |
